# Doubling healthy lifespan using drug synergy

**DOI:** 10.1101/153205

**Authors:** Tesfahun Dessale, Krishna Chaithanya Batchu, Diogo Barardo, Ng Li Fang, Vanessa Yuk Man Lam, Linfan Xiao, Markus R. Wenk, Nicholas S. Tolwinski, Jan Gruber

## Abstract

Pharmacological interventions that target human ageing would extend individual healthspan and result in dramatic economic benefits to rapidly ageing societies worldwide. For such interventions to be contemplated they need to comprise drugs that are efficacious when given to adults and for which extensive human safety data are available. Here we show that dramatic lifespan extension can be achieved in *C.elegans* by targeting multiple, evolutionarily conserved ageing pathways using drugs that are already in human use. By targeting multiple synergistic ageing pathways, we are able to slow ageing rate, double lifespan and improves healthspan while minimize developmental and fitness trade-offs. Moreover, we established that there is no synergistic benefit in a *daf-2* or *daf-7* background, implying the involvement of the TGFβ and IGF pathways in this synergy. Employing lipidomics and transcriptomics analysis we found lipid metabolism to be affected resulting in increased monounsaturated fatty acids (MUFA) and decrease membrane peroxidation index. Our best drug combination showed a conserved lifespan extension in fruit flies. To the best of our knowledge, this is the largest lifespan effect ever reported for any adult-onset drug treatment in *C. elegans*. This drug-repurposing approach, using drugs already approved for humans to target multiple conserved aging pathways simultaneously, could lead to interventions that prevent age-related diseases and overall frailty in a rapidly ageing population.

---

Some of the most important findings in biogerontology are the evolutionarily conserved pathways that regulate lifespan^1-6^. In model organisms, mutations affecting these pathways can often extend lifespan by between 30% and 100%^7,8^. Combining different genetic mutations can result in synergistic lifespan extension^9-11^. By contrast, the effect of pharmacological interventions are typically much weaker, even when targeting the same pathways^2,12,13^, but this genetic synergy suggests a potential strategy for the design of novel pharmacological interventions by simultaneously targeting multiple evolutionarily conserved ageing pathways. To date there is little data on synergistic effects of pharmacological interventions on lifespan^14^. Here we report an *in vivo* approach to design novel multi-drug ageing interventions. We show how multi-drug interventions leverage pathway synergies to maximize effect size, while minimizing side effects and detrimental developmental tradeoffs through targeting distinct but interacting ageing pathways. Our ultimate aim was to design a purely pharmacological, adult-onset intervention with lifespan efficacy similar or better than those of canonical aging mutations did. Due to the lack of generally accepted biological markers for ageing^15-17^, lifespan studies are currently the only way to test the efficacy of ageing interventions^18^. We therefore developed our candidate drug combinations in a short-lived model organism, the nematode *Caenorhabditis elegans*. Using our approach, we identify two triple drug combinations that extend lifespan and healthspan to an extent greater than any previously reported pharmacological intervention in *C. elegans*^13^. Our synergistic drug combinations show effect sizes comparable to the classical ageing mutations while avoiding most of the tradeoffs associated with them^19-21^. We find that these interventions actually increase some markers of performance while slowing biological ageing rate. Moreover, we identified TGFβ as a key contributor and required pathway in mediating these synergistic effects. We also find that worms treated with synergistic drug combinations have higher MUFA:PUFA ratios and a decrease in membrane lipid peroxidation index. Finally, we confirm that this synergistic effect is also present in the fruit fly *Drosophila melanogaster*.

## Results

### Pathway and compound selection in *C. elegans*

Based on existing literature, we identified a set of well-characterized and evolutionarily conserved ageing pathways and lifespan extension mechanisms (Supplementary Table S1). We chose to target AMP activated protein kinase (AMPK), mammalian target of rapamycin (mTOR), caloric restriction (CR), C-Jun N-terminal kinases (JNK) and mitohormesis/mitochondrial metabolism as primary longevity regulatory pathways. For each pathway, we identified drugs and drug-like molecules reported to extend lifespan in at least one common model organism (nematodes, fruit flies or mice). We were interested in drugs that might eventually be tested in humans so we favored drugs with reported efficacy in mammals or that are already approved for human use. Based on these criteria, we initially selected eleven candidate drugs (Supplementary Table S1). We added allantoin to our study based on a report showing lifespan effects in *C. elegans* and a transcriptional analysis suggesting that its mode of action is unusually distinct from other compounds^22^. In order to test the lifespan effects, which can be highly sensitive to details in experimental conditions and can be variable between different laboratories^19,23^, we first carried out operator-blinded confirmatory lifespan studies using the previously reported optimal dose for each drug. We found that only five compounds extended lifespan reproducibly under conditions used in our laboratory (Fig. 1, Extended data Fig. 1, Supplementary Table S2). Generally, lifespan extension effects in our hands tended to be smaller compared to previous reports^7,22,24-26^ (Fig. 1, Extended data Fig. 1, Supplementary Table S2).

**Figure 1.**
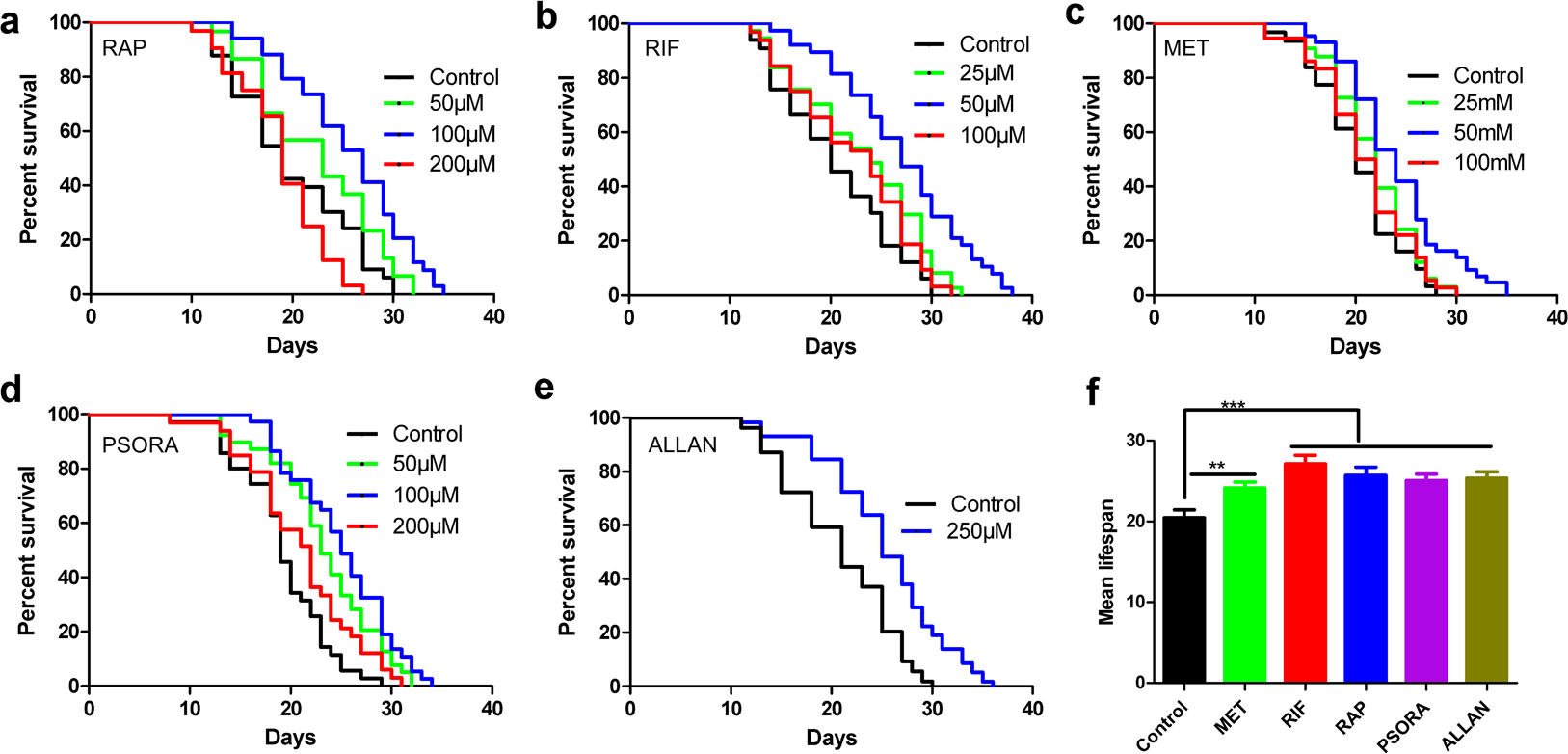
Single drugs extend lifespan of wild type *C. elegans*. Wild type N2 worms treated with different doses of **a**, RAP **b**, RIF **c**, MET **d**, PSORA and **e**, ALLAN have longer lifespan. **f**, Mean lifespan of worms treated with the optimal dose of each drug, mean ± SD. Each drug treatments resulted in a statistically significant lifespan extension at their optimal dose. **P < 0.001, ***P < 0.0001, log-rank with adjustment for multiple comparisons.

### Gene expression signature of single drug treatments

To examine drug modes of action, we carried out transcriptomics analysis and determined differentially expressed genes (DEG) and pathway enrichments relative to untreated controls. Rifampicin (RIF) and rapamycin (RAP) followed by psora-4 (PSORA) caused the most extensive changes in gene expression, both at the gene and pathway level, while metformin (MET) and allantoin (ALLAN) showed comparatively smaller effects on gene expression (Fig. 2a Supplementary Table S3). While transcriptional analyses revealed significant overlap between affected genes, we also identified sets of genes that were unique to each individual compound (Fig. 2a). MET shows significant overlap with the other drugs, having the smallest unique set. Almost all genes affected by MET and PSORA were also affected by at least one other drug in the set (Fig. 2a). ALLAN causes fewer gene-expression changes overall than any of the other compounds, but 48% of the genes affected by ALLAN are unique, that is, not affected by any of the other drugs, while only 17% of the genes affected by MET are unique (Fig. 2a). Globally, effects of RIF and RAP were quite different from each other and from any of the other drugs. We confirmed this by principle component analysis (PCA) which showed that RIF, RAP and PSORA are well separated by the first three principle components. ALLAN and MET, by contrast, are more variable and very close to each other and to untreated control (Fig. 2b, Extended data Fig. 4c).

**Figure 2.**
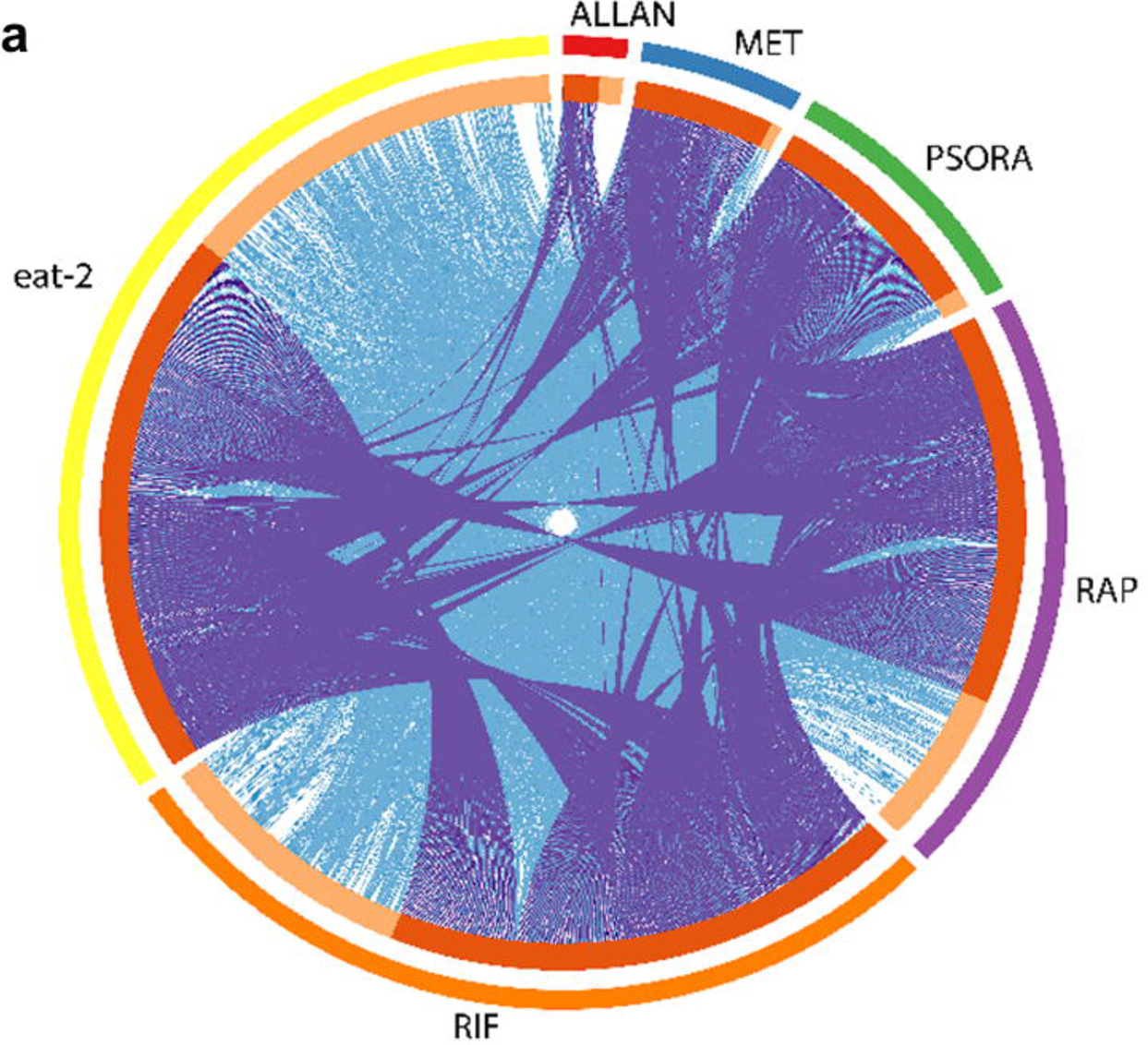
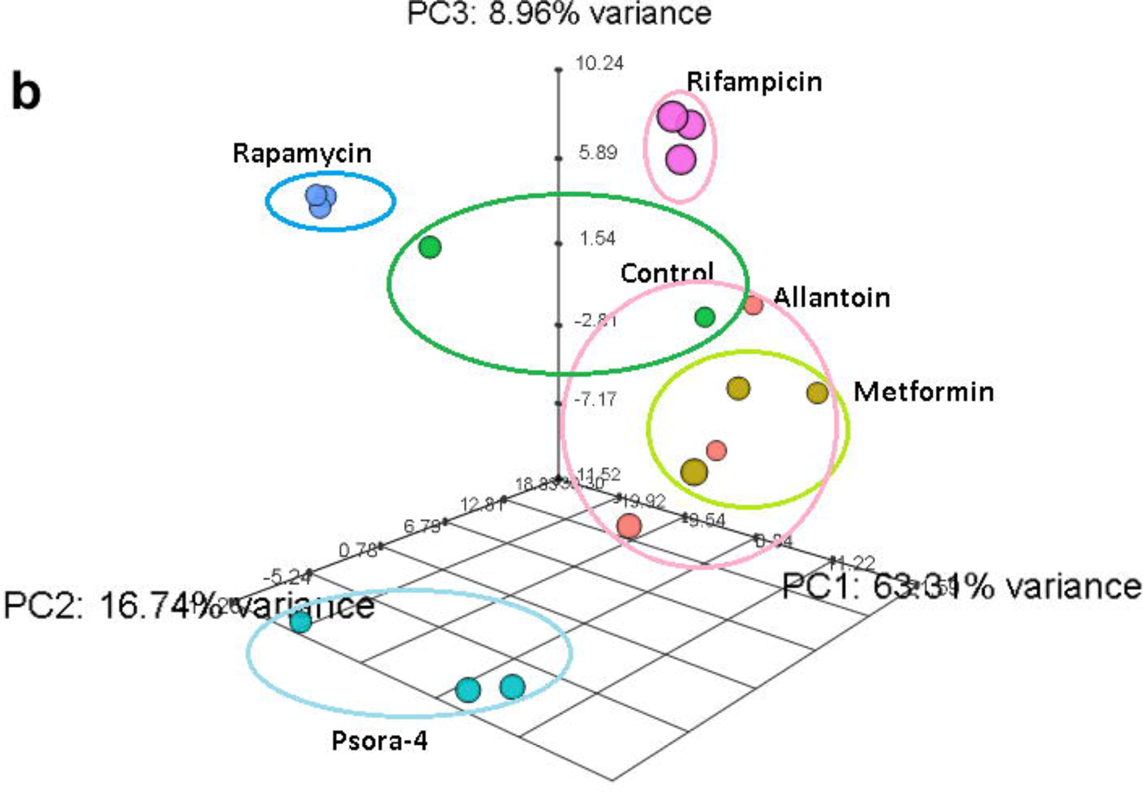
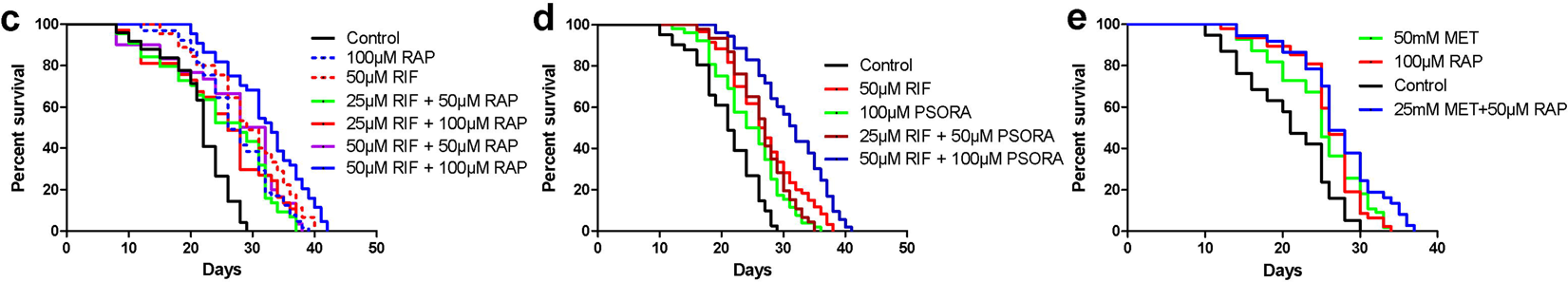
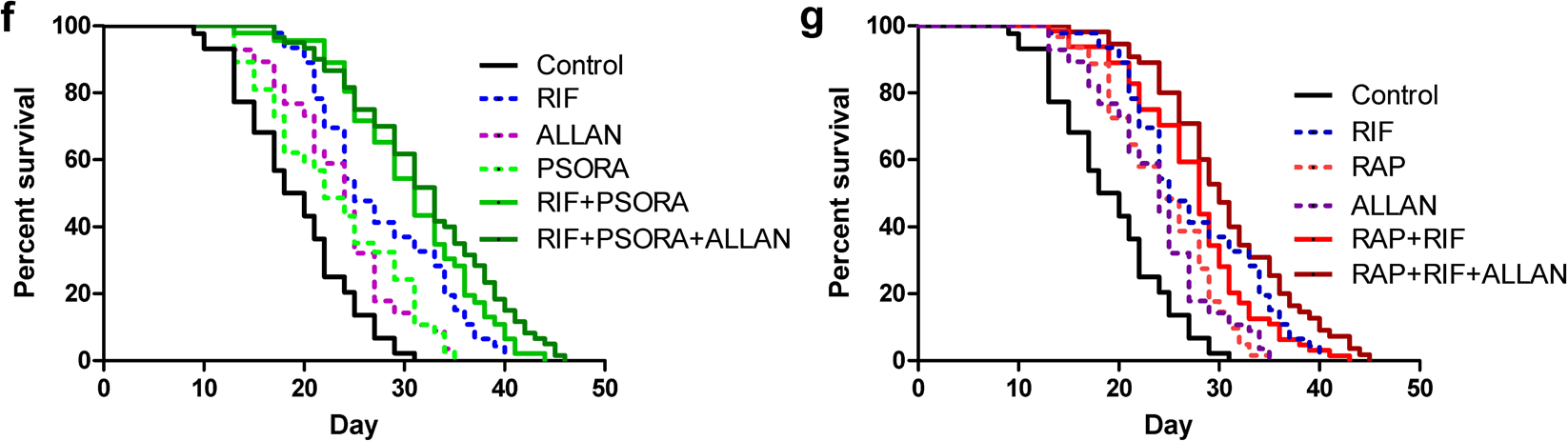
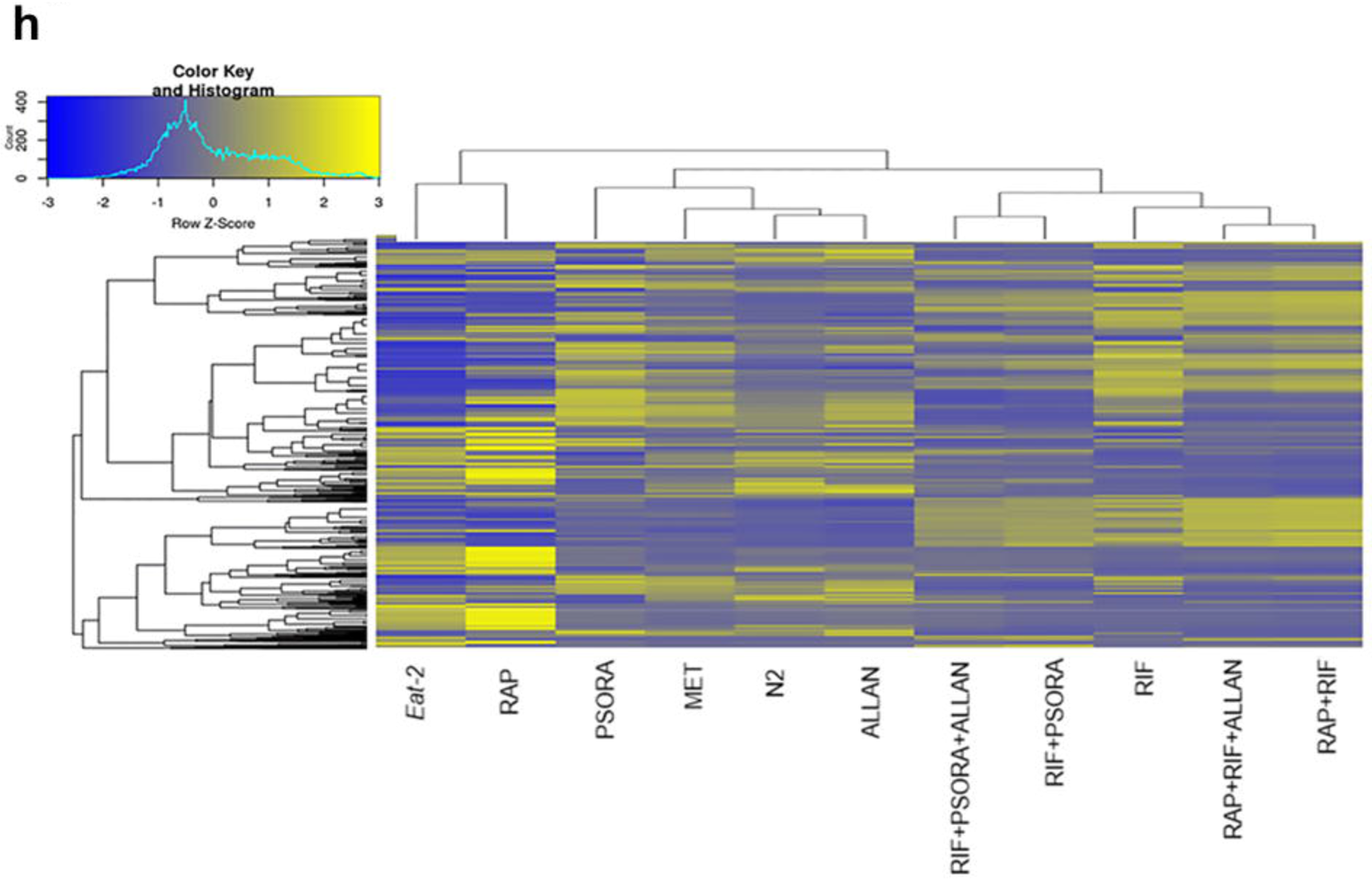
Double and triple drug combinations showed synergistic lifespan extension; **a,** Circos plot illustrating gene overlap for single drugs and *eat-2* mutant transcriptome. Purple lines link genes shared by multiple drugs. Blue lines link different genes which fall into the same GO term. A greater number of purple and blue links and longer dark orange arcs indicates greater overlap among the DGE and GO terms of each drug. (Minimum overlap = 3, minimum enrichment = 1.5, P value < 0.01). **b,** Three dimensional principle component analysis based on DGE (PCA). **c,** RAP+RIF and **d,** RIF+PSORA results in synergistic lifespan extension (P<0.0001, log-rank with adjustment for multiple comparisons). **e,** RAP+MET did not show further mean lifespan extension but showed further maximum lifespan extension compared to single drug treatments. **f**, RIF+PSORA+ALLAN and **g.** RAP+RIF+ALLAN showed synergistic lifespan extension (P<0.0001, log-rank with adjustment for multiple comparisons). **h**, Heatmap showing hierarchical clustering (distance metrics and linkage algorithms) of single drugs and synergistic combinations. n = 2000 for RNAseq and 50-100 for lifespan studies. *P<0.01, **P<0.001, ***P<0.0001

### Drug synergies and lifespan extension

We next determine effect of different drug combinations on lifespan in nematodes. We began with MET and RAP due to their translational importance in humans^14,27^. However, RAP combined with MET at their optimal doses did not show any further benefit on lifespan (Extended data Fig. 2j,p). We carried out additional trials, exploring all possible combinations of optimal and half-optimal dose for MET and RAP (Extended data Fig. 2j-p). Combination of RAP and MET each at their half-optimal dose result in further extension in maximum lifespan (Fig. 2e, Extended data Fig. 2m,s) but did not result in further mean lifespan extension (Extended data 2p). These weak benefits of combining MET with RAP are consistent with results in mice where the existing evidence suggests that addition of MET to RAP results in some additional benefits mainly in males with only weak (if any) additional benefits in females^14^.

We then systematically determined the efficacy of all ten possible pairs out of the five candidate drugs (Extended data Fig. 2a-i). For these trials we define synergy based on the Higher Single Activity (HSA) model^28^; Drug combinations are considered synergistic if the combinatorial effect is significantly larger than any of the combination’s single drug effect at the same (ideal) concentrations as in the combination. Using this definition, we identified two synergistic combinations out of the ten pairs tested (Fig. 2c, d, Extended data Fig. 2 and Supplementary Table S2). The synergistic pairs comprise RIF with either RAP or PSORA. This means that two out of the three pairs formed by the drugs that are most well-separated in the PCA are synergistic (Fig. 2b, Extended data Fig. 4a,b). Of the remaining eight combinations, four extended lifespan as much as their best component while the other four combinations nullified each other and did not result in lifespan extension and none were toxic (Extended data Fig. 2, Supplementary Table S2).

Lifespan extension resulting from the two synergistic pairs, while larger than those of previously published drug effects, were still smaller than the benefits seen with genetic mutations^9^. This raises the question whether the effect size seen with the synergistic pairs represented the maximum achievable by adult-onset drug treatment or if further benefits could be gained by adding additional compounds. Since exhaustive combinatorial search of all thirty possible triple drug combinations was not feasible, we explored selected triple combinations based on the original synergistic pairs and taking into account our analysis of transcriptional changes for the single drugs. We first evaluated the combination of the three drugs best separated by PCA and involved in the two initial synergistic pairs (RAP, RIF and PSORA). However, nematodes treated with this triple combination had a shorter lifespan than seen with either of the dual combinations (Extended data Fig. 3a Supplementary Table S2). We then tested three drug combinations including ALLAN as the third component. As explained above, ALLAN has a gene expression signature that is quite distinct from the rest of our drugs. Furthermore, unlike for MET, the unique gene set of ALLAN (light orange arc in Fig. 2a) also shows no overlap at the level of GO terms (blue links) with any of the other drugs or with *eat-2*. This further supports the notion that ALLAN has a unique mode of action^22^. Therefore, we tested addition of ALLAN to the two synergistic double combinations. This addition resulted in significant additional mean and maximum lifespan extension in both cases (Figures 2f,g, Extended data Fig. 3d-f and Supplementary Table S2). The best triple combination (RAP, RIF, ALLAN) doubles mean lifespan, resulting in a median lifespan of up to 44 days and a maximum lifespan of 50 days (Supplementary Table S2). This effect size is comparable to the canonical ageing mutations and is, to the best of our knowledge, the largest lifespan extension ever reported in *C. elegans* with any adult-onset drug intervention^29^. By contrast, addition of ALLAN to the non-synergistic combination (RAP+PSORA) did not show any further benefit (Extended data Fig. 3c). Finally, to test whether, despite its high overlap in terms of gene expression changes, addition of MET would results in any further lifespan benefit on dual combinations, we add MET as a third component to one of the synergistic dual combinations RAP+RIF. However this triple combination (RAP+RIF+MET) resulted in toxicity (Extended data Fig. 3b).

**Figure 3.**
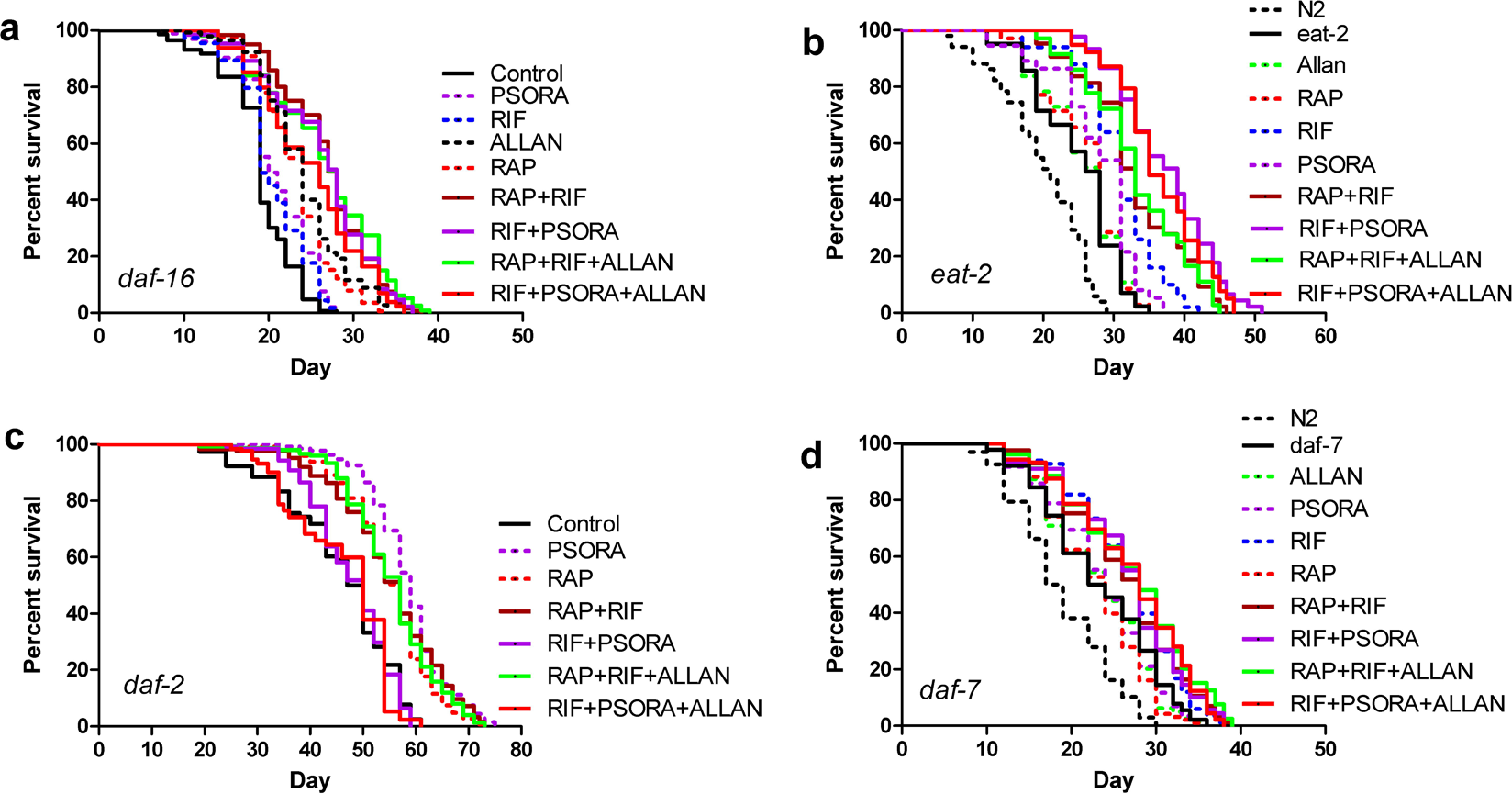
Drug combinations did not show synergy on *daf-7*, *daf-2* and *daf-16* mutants. **a,** all dual and triple synergistic combinations resulted in lifespan extension in *daf-16* mutants but effect size does not exceed individual drug effects. **b,** Of the single drugs only RIF extends lifespan of *eat-2* (P<0.05, log-rank with adjustment for multiple comparisons). RIF+PSORA resulted in synergistic lifespan extension in *eat-2* mutants (P<0.05, log-rank with adjustment for multiple comparisons). **c,** None of the synergistic combinations showed synergy in *daf-2* mutants. **d,** RIF alone extends lifespan of *daf-7* mutants (P<0.05, log-rank adjustment for multiple comparisons) but combinations fail to result in further synergistic lifespan extension. At least 50 worms per condition.

### Mechanisms of drug synergy

*Daf-16*/FOXO is a transcription factor that plays a key role in lifespan determination in model organisms and likely also in humans^30,31^. To explore the mode of action of our synergistic drug combinations we determined their dependence on *daf-16* in mutant worms deficient in this pathway. Individually, RAP and ALLAN lifespan extension was *daf-16*-independent while RIF lifespan extension was fully and PSORA was partially *daf-16* dependent (Fig. 3a, Extended data Fig. 8). The synergistic combination RAP+RIF comprising one *daf-16* dependent (RIF) and one *daf-16* independent drug (RAP), and even the RIF+PSORA combination of two *daf-16* dependent drugs, still showed synergistic lifespan extension in *daf-16* mutants (Fig. 3a, Extended data Fig. 8). This surprising data suggest that synergies can be *daf-16* independent, even if individual constituent drugs are *daf-16* dependent.

Several of the drugs tested have previously been reported to be CR mimetics^17,32-34^. We therefore tested each synergistic combination and its constituent compounds in a *C. elegans* model of CR (*eat-2* mutants). We found that only RIF among the single drug treatments further extended lifespan in *eat-2* mutants (Fig 3b, Extended data Fig. 8), suggesting that all except RIF may function as CR mimetic in this sense. The synergistic combinations both comprise one CR mimetic with one non-CR mimetic (RAP+RIF and RIF+PSORA). Only RIF+PSORA resulted in further lifespan extension and addition of ALLAN, a second CR mimetic, did not result in further benefits.

To further explore the mechanism of drug synergy we determine the transcriptome profile for each synergistic combination and selected non-synergistic drug combinations. Transcriptomic analysis in worms treated with single, double and triple drug exposure for the synergistic and non-synergistic combinations revealed that the transcriptome of CR mimetic single drugs is clustered together with *eat-2*, whereas the non-CR mimetic RIF is different from *eat-2* and from all other single drugs (Fig. 2h). Interestingly, the transcriptomes of synergistic combinations are clustered together and the pattern is different from those of their constituent single drugs and from that of *eat-2* mutants (Fig. 2h, Extended data Fig. 4c,d). Moreover, we found that only on pathway, transforming growth factor beta (TGFβ), was commonly enriched in all synergistic dual and triple combinations (Fig. 4a, Extended data Fig. 6c, 7a, Supplementary Table S4) relative to untreated control. To identify pathways that were further enriched by the drug combinations relative to their constituent single drugs, we used the single drug transcriptome as background control (see methods) for pathway analysis. Using this approach, we found that all synergistic combinations, again, affected TGFβ amongst the pathways differentially impacted relative to their parent drugs while none of the non-synergistic combinations did so (Supplementary Table S5). Because some drug combinations resulted in further lifespan extension in *eat-2* mutants relative to their constituent drugs, we tested whether TGFβ was enriched in the transcriptomes of N2 animals treated with synergistic drug combinations relative to untreated *eat-2* worms. Again, TGFβ was the only pathway enriched in all synergistic combinations (Extended data Fig. 7b Supplementary Table S6). Overall, we found that the TGFβ signaling pathway was commonly and exclusively enriched in synergistic combinations (Fig. 4b, Extended data Fig. 7a,c, Supplementary Table S7).

**Figure 4.**
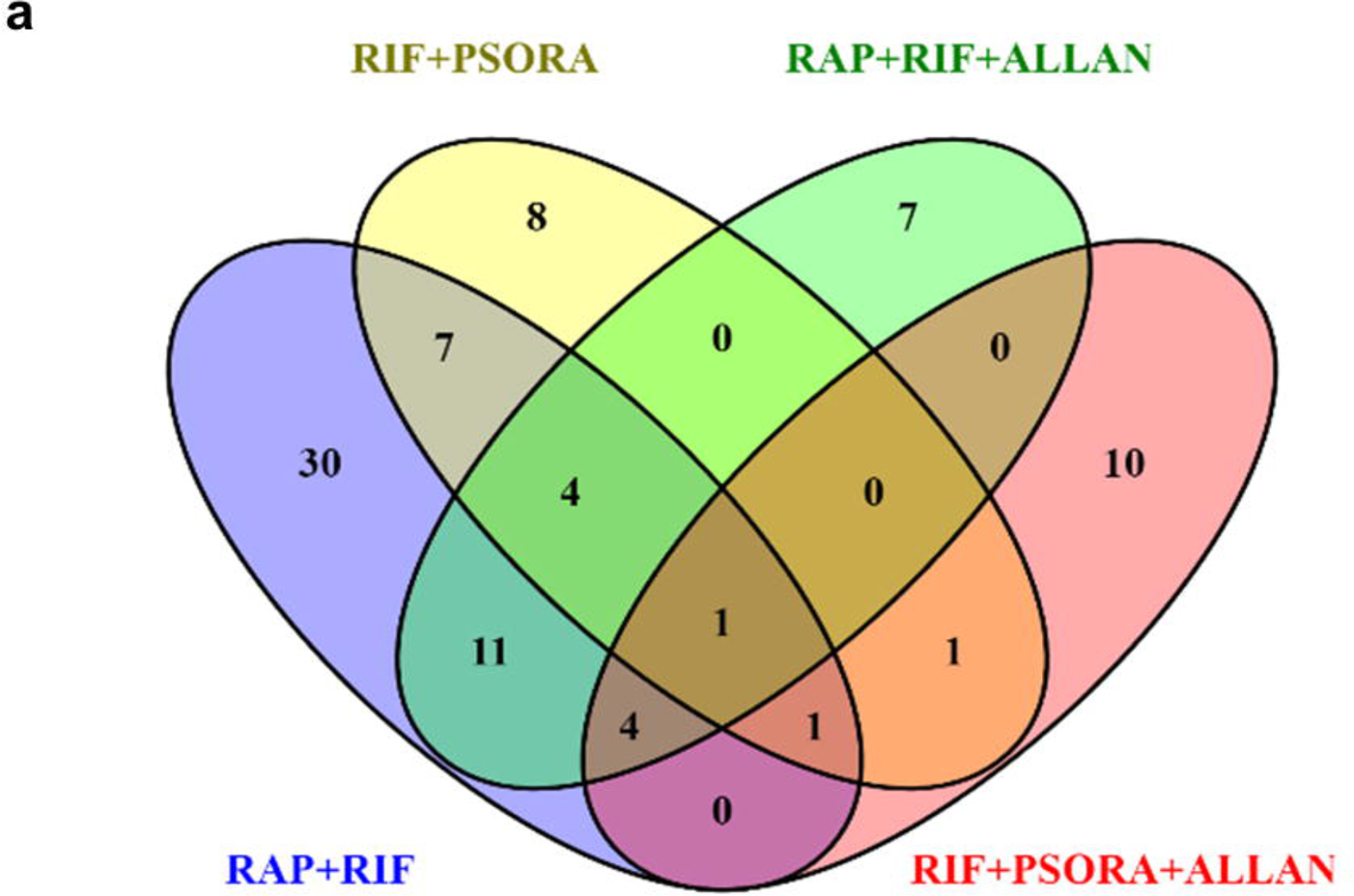
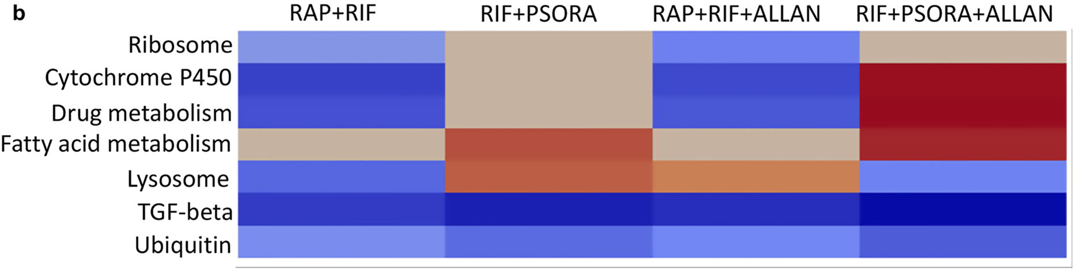
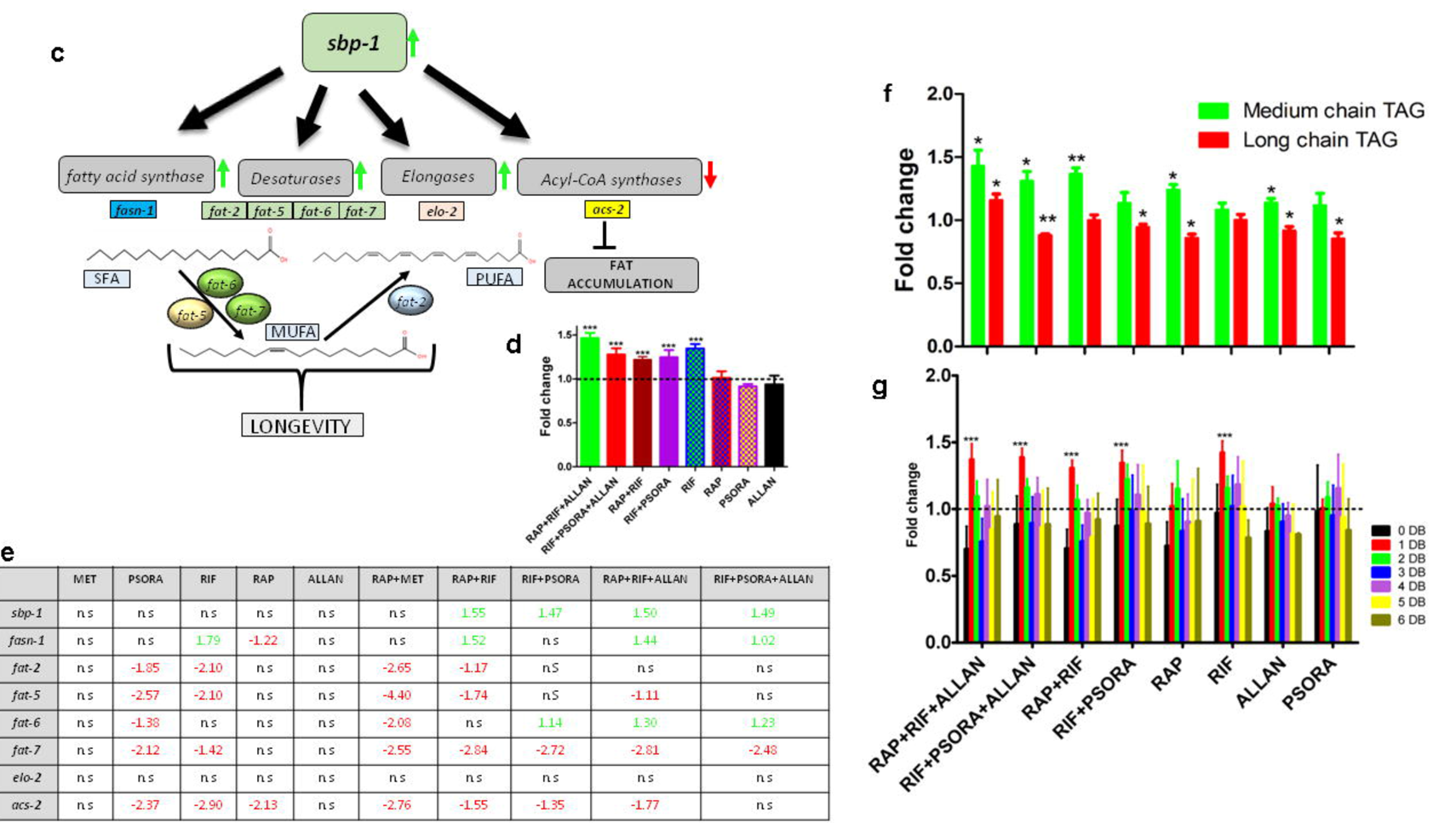
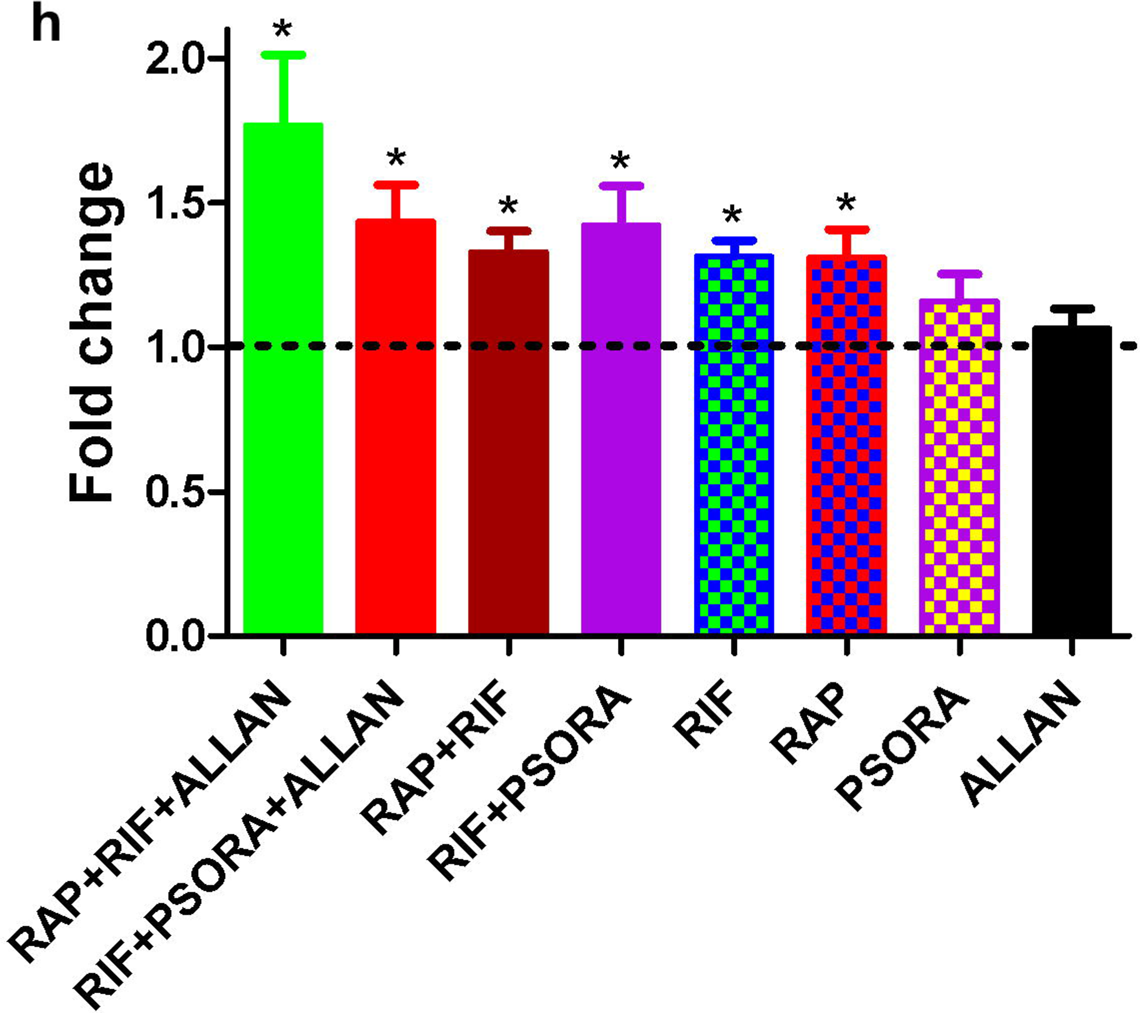
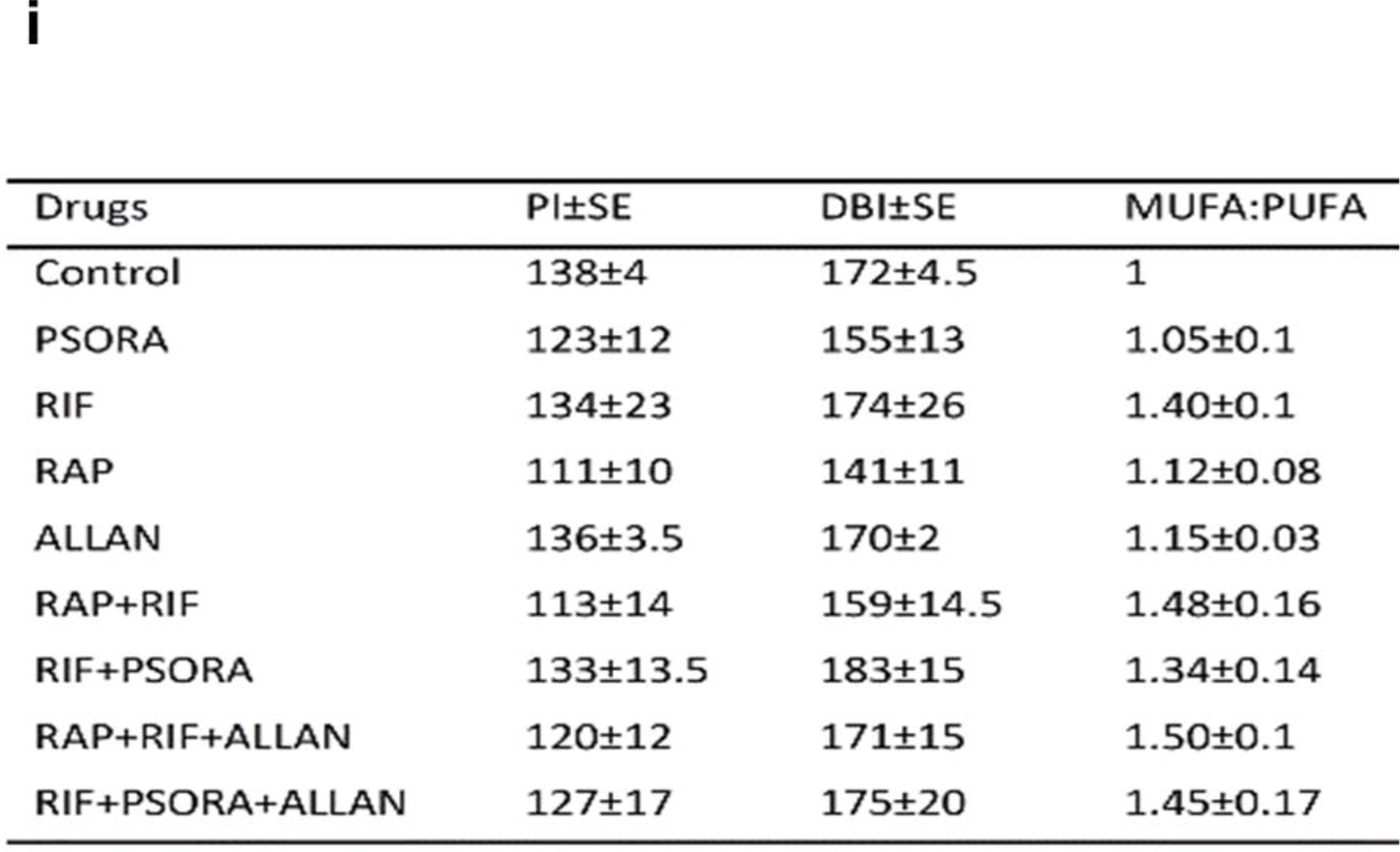
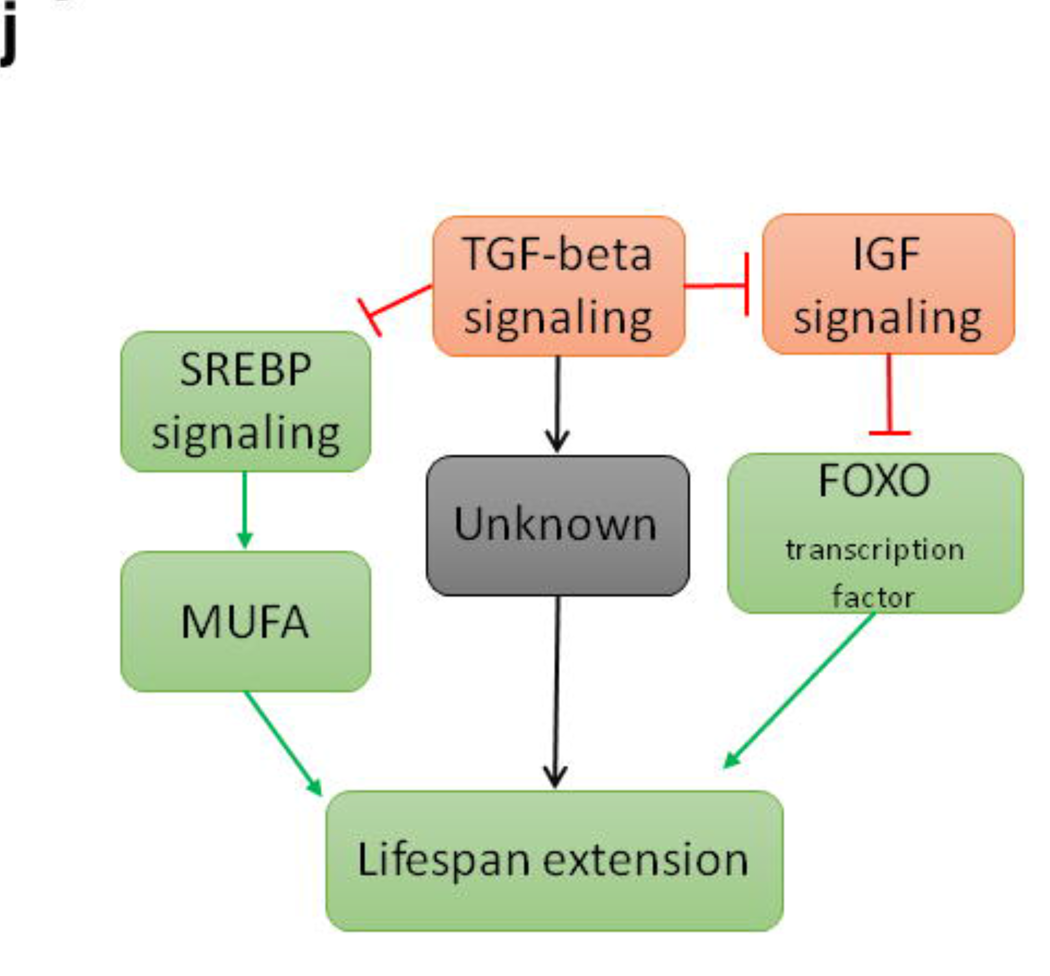
Transcriptomics and lipidomics profiles explain the mechanism of drug synergy. **a,** Venn diagram for pathways enriched by synergistic drug combinations. Only is commonly enriched in all synergistic combinations. **b,** Heatmap of pathways enriched by synergistic drug combinations compared to their constituent single drugs. Only is commonly enriched in all synergistic combinations. **c,** Mechanism of MUFA containing lipid accumulation and lifespan extension. **d,** Total triacylglycerol content normalized to the control, three biological replicates, 2500 worms per condition, (Mean ± SD, one-way ANOVA) e, Fat metabolism-related gene expression profile, log fold change, P value < 0.05. **f,** TAGs categorized as those containing medium chained and long chained acyl chains, normalized to control, three biological replicates, 2500 worms per condition (Mean ± SD, one-way ANOVA). **g,** Phosphatidylcholine species abundance based on double bond, three biological replicates, 2500 worms per condition (Mean ± SD, one-way ANOVA). **h,** Total sphingomyelin content normalized to the control, three biological replicates, 2500 worms per condition, (Mean ± SD, one-way ANOVA). **i,** Double bond index, per oxidation index and MUFA/PUFA ratios calculated for phosphatidylcholine species for all conditions, MUFA/PUFA is normalized to the control. Three biological replicates, 2500 worms per condition (Mean ± SD), **j.** Hypothesized mechanism of lifespan extension by drug synergies. Red–inhibition, green–activation, black–unknown. *P<0.01, **P<0.001, ***P<0.0001.

### TGFβ is required for drug synergy

The specific and consistent enrichment of TGFβ suggests that TGFβ may play a role in mediating the observed synergistic lifespan extension. Previously it has been shown that TGFβ (*daf-7*) mutation extends lifespan *via* insulin signaling and that both *daf-2* and *daf-7* regulate the transcription of *daf-16* dependent genes^35^ (Extended data Fig. 7d). To test the role of the IGF pathway in this effect and to determine if the lifespan of already long-lived *daf-2* mutants could be further extended using our drug combinations, we first determined lifespan of *daf-2* mutants for each combination. Only the RAP-based combinations, RAP+RIF and RAP+RIF+ALLAN, showed lifespan extension in *daf-2* mutants. However, the effect size was similar to that of RAP alone and none of the combinations resulted in further synergy (Fig. 3c, Extended data Fig. 8, Supplementary Table S2). PSORA treatment alone extended the lifespan of *daf-2* mutants, but none of the PSORA based combinations, RIF+PSORA and RIF+PSORA+ALLAN, extended lifespan in *daf-2* mutants (Fig. 3c and Extended data Fig. 8). To test our hypothesis that TGFβ was involved in mediating drug synergies, we next tested the efficacy of all synergistic combinations and their components in *daf-7* mutants. None of the combinations show synergistic effects in the *daf-7* mutants, suggesting that synergy requires *daf-7*, even though some of the drugs singularly still extend lifespan (Fig. 3d, Extended data Fig. 8).

### Transcriptome, Lipidome and Drug synergy

Long-lived mutants such as *age-1* and *daf-2* have previously been shown to exhibit metabolic perturbations resulting in increased production and storage of fats^36^. Furthermore, TGFβ/*daf-7* regulates triacylglycerol (TAG) metabolism^37^ and *daf-7* mutants are known to store more fats^36^. These links in conjunction with our transcriptomics data led us to investigate whether treatment with synergistic drug combinations resulted in modifications in lipid profiles consistent with TGF inhibition. (Supplementary Table S3). First we explored the transcriptomics data for changes that might affect lipid composition and found that synergistic drug combinations indeed resulted in a significant up regulation of the *C. elegans* SREBP-1c homolog - *sbp-1*, a master transcription factor controlling several lipogenic genes, and genes coding for “desaturases” responsible for MUFA synthesis^38-40^ (Fig. 4c,e). We then employed an MS-based lipidomics assay to determine changes in lipid compositions that might result from these drug treatments. Worms that were exposed to the synergistic drug combinations have a significant rise in TAG reserves (Fig. 4d), with increased abundances in those TAG species that contained the medium chained saturated fatty acids (Fig. 4f). In addition, we also observed a marked increase in the MUFA:PUFA ratios (Fig 4g, Extended data Fig. 7e). Because of these changes, there was a significant decline in lipid peroxidation index (PI) (Fig. 4i). A low PI, indicates that lipids contain fewer carbon-carbon double bonds, making them less susceptible to peroxidation and this has been previously associated with increased lifespan, probably related to lower susceptibility to lipid peroxidation^41,42,43^. A reduction in the susceptibility to lipid peroxidation suggests better resistance to oxidative stress^44,45^ and worms treated with synergistic drug combinations indeed showed such an increase in resistance (Fig. 5f). We also observed an increase in total sphingomyelin (Fig. 4h); an event that has previously shown to elicit autophagy-dependent lifespan extension in the nematodes^46^.

**Figure 5.**
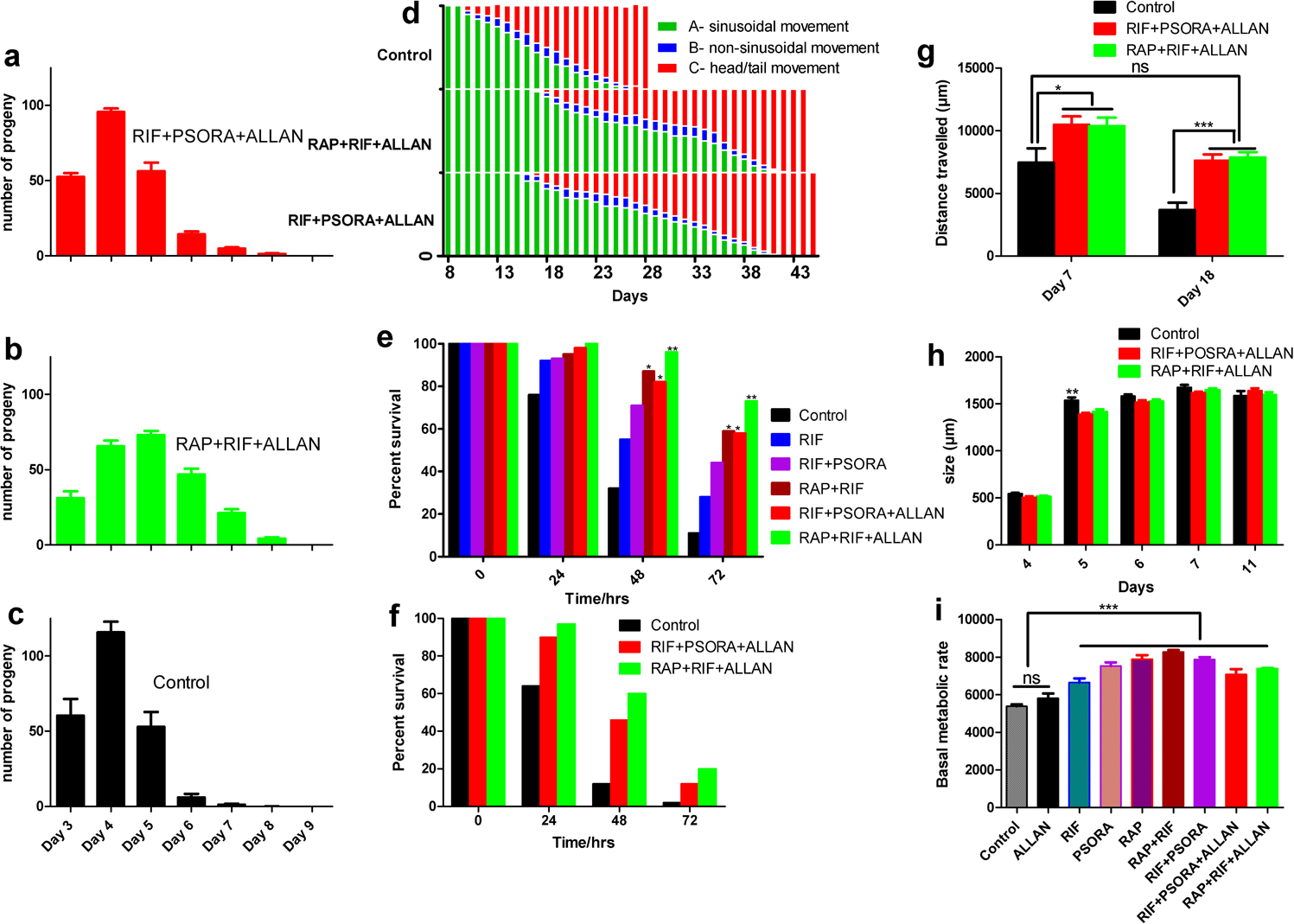
RIF+PSORA+ALLAN and RAP+RIF+ALLAN improve healthspan without tradeoffs. **a-c,** RIF+PSORA+ALLAN and RAP+RIF+ALLAN shows extension of reproductive span but no effect on total fertility, n=10 worms per condition. **d,** health span, n>150 worms per condition, P<0.0001, one-way ANOVA. **e,** resistance to heat stress, n=50 worms per condition. **f,** resistance to paraquat stress, n=50 worms per condition. **g,** distance travelled n=50 worms per condition, one-way ANOVA. **h,** development measured by size of worms n=10 worms per condition. **i,** basal metabolic rate, n=80 worms per condition, one-way ANOVA, Bonferroni multiple comparison. *P<0.01, **P<0.001, ***P<0.0001

### Drug synergy improves health span and delays ageing

Suppression of energy metabolism and inhibition of the electron transport chain (ETC) can extend lifespan in *C. elegans*^47,48^. In our experiments, however, drug treatments did not have inhibitory effects on basal metabolic rate, instead several of this drugs and drug combinations increased basal respiration (Fig. 5i), excluding ETC inhibition as explanation for the observed lifespan effects. The most common tradeoffs related with lifespan extension are developmental delay, slower growth rate and reduced fecundity^49,50,51^. We found that in worms treated with either of the synergistic drug combinations, there is evidence for a delay in peak egg laying and an extension of reproductive span but no change in total fertility. (Fig. 5a-c, Extended data Fig. 9a).

Next, we assessed several parameters of fitness, stress resistance and health: speed of spontaneous movement, tolerance to heat shock and oxidative stress resistance and a motility based health span score based on the scoring scheme of Herndon *et al*^52^. Control animals spent about 50% of their lifespan in the optimal (best) health category while nematodes exposed to the RIF+PSORA+ALLAN and RAP+RIF+ALLAN spent 57% and 63%, respectively, of their already extended lifespan in optimal health (Fig. 5d, Extended data 9c). This means that their health span was significantly extended both in absolute and relative terms. Indeed, more than half of the treated animals were still in optimal health even after the last control animal had died (Fig. 5d, Extended data Fig. 9c). Surprisingly, animals treated with RAP+RIF+ALLAN and RIF+PSORA+ALLAN also performed significantly better in the spontaneous movement assay than control animals at most ages. In fact, old treated animals (day 18 of age) showed performance in this assay that was indistinguishable from young (day 7) controls (Fig. 5g). Moreover, treated animals experienced improvements in resistance to heat and oxidative stress (paraquat) assays, (Figure 5e,f, Extended data Fig. 9b). These observations indicate that animals treated with synergistic drug combinations not only enjoyed greater lifespan with limited detected tradeoffs, but also improved health span, increased stress resistance, and even increased physical performance. Such observations raise the important question if these lifespan benefits are due to an actual delay in biological ageing rate. The rate of ageing can be expressed as mortality rate doubling time (MRDT). We determined MRDT of animals exposed to synergistic drug combinations using Survcurv^53^, and found MRDT to be significantly longer in RIF+PSORA+ALLAN treated animals compared to controls (MRDT of control = 3 days and RIF+PSORA+ALLAN = 3.7 days, P value < 0.0001). The initial mortality rate was significantly lower for both synergistic combinations ((IMR of control = 2.7e^-3^, IMR of RIF+PSORA+ALLAN = 8.5e^-4^, RAP+RIF+ALLAN = 9.3e^-4^, P value < 0.001). This means that the RIF+PSORA+ALLAN drug combinations extend lifespan by making worms age more slowly while at the same time making them move faster/increasing spontaneous activity and increasing their resistance to stress when young. (Extended data Fig. 10).

### Drug synergy is conserved in Drosophila

One of the key questions for ageing studies is whether lifespan effects are conserved across species. We therefore tested whether the synergistic lifespan extensions we found in *C. elegans* are conserved in *Drosophila melanogaster*. RAP has previously been shown to extend lifespan of fruit flies^54^ while MET has been shown to be ineffective^55^. We systematically tested all single drugs and all synergistic combinations in male fruit flies. First, we showed that RAP and ALLAN extend mean lifespan whereas RIF and PSORA extend only the maximum lifespan of flies (Fig. 6 Supplementary Table S2). However, the RAP based combinations RAP+RIF and RAP+RIF+ALLAN showed conserved synergistic lifespan extension in flies (Fig. 6a). The PSORA based combinations; RIF+PSORA and RIF+PSORA+ALLAN did not show further mean lifespan extension (Fig. 6b). However, RIF+PSORA showed synergistic maximum lifespan extension compared to single drugs (Fig. 6c). Comparison of the Gompertz parameters of the mortality trajectories (Extended data Fig. 10b) showed that the baseline mortality rate was not significantly different in drug combination treated flies (IMR difference of RAP+RIF+ALLAN = 2.9e^-1^, RIF+PSORA+ALLAN = 4.8e^-1^, NS) compared to untreated control. However, the rate of increase of mortality with age is significantly lower in drug combination treatments (Gompertz different b of RAP+RIF+ALLAN = 2.20e^-4^, P-value < 0.001, Gompertz different b of RIF+PSORA+ALLAN = 1.15e^-2^, P-value <0.05) relative to the control. It should be noted that cohort size of flies was smaller than for worms and that we did not re-optimize drugs for the fly experiments. However, these data confirms that some of the detected synergies appear to be evolutionary conserved.

**Figure 6.**
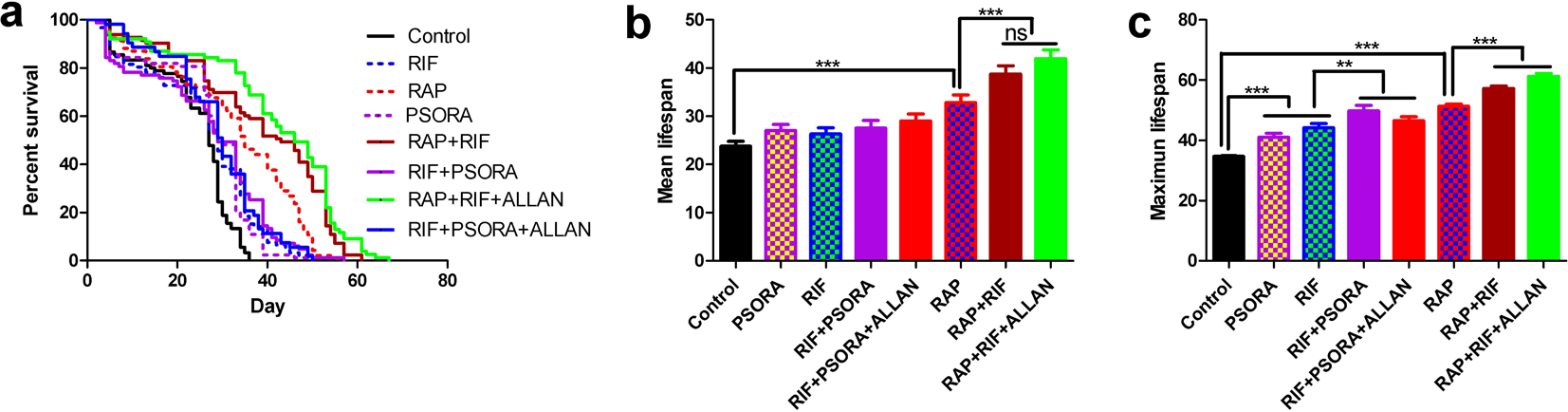
Drug synergies resulted in conserved lifespan extension in *Drosophila melanogaster*. **a,b,c** RAP extends both mean and maximum lifespan. RAP+RIF and RAP+RIF+ALLAN resulted in synergistic mean and maximum lifespan extension. RIF and PSORA did not extend mean lifespan of flies. Their combination also did not extend mean lifespan. However, RIF and PSORA extends maximum lifespan significantly. RIF+PSORA and RIF+PSORA+ALLAN resulted in further maximum lifespan extension, close to additive (P value < 0.001) but not mean lifespan. n = 80 flies per condition. One-way ANOVA, Bonferroni multiple comparison for mean and maximum lifespan and Log-rank test for survival analysis. *P<0.01, **P<0.001, ***P<0.0001

## Discussion

By targeting multiple overlapping ageing and longevity-related pathways we succeeded in designing synergistic pharmacological interventions that, even when only applied to animals from adult age, more than doubled healthy lifespan in *C. elegans*. This effect size is comparable to that of the classical ageing mutations and, to the best of our knowledge, is the largest reported for any adult-onset pharmacological intervention^13^. However, unlike in ageing mutations we detect very limited detrimental evolutionary or fitness tradeoffs associated with this lifespan extension by drug synergies ^19^. In particular, we did not detect any decrease in oxygen consumption or total fecundity. The latter result is consistent with a previous study showing benefits of an adult-onset intervention using resveratrol without reduced fecundity^56^. These data suggest that there is significant post-developmental plasticity of lifespan that can be exploited using adult-onset pharmacological interventions. In fact, treated animals outperform age-matched controls in some performance assays and old treated animals have the physical appearance and performance comparable to much younger controls (Fig. 5g). Mortality analysis suggests that one synergistic drug treatment slows basic biological ageing rate by ~20% (Extended data Fig. 10). To test for evolutionary conservation, we determined lifespan benefits in the fruit fly *Drosophila melanogaster*, confirming qualitative conservation for two of the four synergistic drug combinations. Nematodes are evolutionarily more distant from fruit flies than they are from mammals^57^, making mechanisms and pathways conserved between flies and nematodes ancient, and tracing this synergy back to a common ancestor of all three clades.

To explore the mechanism of these synergistic lifespan benefits, we carried out epistasis experiments and transcriptomic analysis, finding that *daf-2/daf-16* (IGF/FOXO) as well as the TGFβ pathway (*daf-7*) are involved in synergistic lifespan extension (Fig. 4j). The connection between IGF/FOXO and TGF is consistent with previous results where *daf-7* regulates lifespan *via* insulin signaling^35^, neuronal TGFβ links food availability and longevity^58^, and *daf-7* regulates metabolism in *C. elegans* and controls fat accumulation^36^. Several previous studies have demonstrated that lifespan extending mutations in *C. elegans* result in alterations in lipid metabolism causing an increase in the MUFA and reduction in PUFA levels^59,60^. In agreement with these reports, our drug synergies resulted in an up regulation of *sbp-1*, a master transcription factor controlling several genes related to lipid metabolism and MUFA synthesis^38-40^ (Fig. 4c,e, Extended data Fig. 5). MS-based lipidomics analysis of nematodes exposed to synergistic drug combinations revealed major changes consistent with activation of *sbp-1*, including accumulation of TAGs and increases in the MUFA:PUFA ratios (both in PC and PE classes). Previously, it has been shown that an increased TAG abundance prolongs lifespan in budding yeast^61^ and *C. elegans*^36,38^. Furthermore, upon closer inspection of the fatty acyl chains bound to the TAG species, we found a significant increase in those that contained medium chained saturated fatty acyls (MCFA) (14:0, 16:0), (Fig. 4f). The findings from lipidomics therefore are consistent with the observed induction of the *sbp-1* and its target lipogenic genes (Figure 4c). Increased abundance of such medium chained saturated fatty acyls has also been reported to be positively correlated with longevity in *C. elegans* previously^59^. Similarly, high abundance of MUFAs has been shown to be associated with extended lifespan in nematodes^62^. In humans, high ratios of MUFA:PUFA have also been found in erythrocyte membranes of children of nonagenarians^63,64^. There has been much interest in the potential for MET and possibly RAP to delay age-dependent decline and disease in humans. Given this translational interest, the limited benefit of combining MET with RAP in both *C. elegans* and mice is somewhat disappointing. However, as our results illustrate, lifespan is determined through complex and interactive biochemical and gene regulatory networks. Intervening simultaneously at multiple points of these networks can result in significant and sometimes surprising benefits^28^. Our proof-of-principle study suggests that beneficial synergistic and additive interactions affecting key longevity pathways are unexpectedly common and evolutionarily conserved. While, to date, ours is the largest systematic transcriptional screen of drug synergy interactions between compounds targeting ageing pathways, it is important to note that this dataset is too small to derive definitive rules by which to detect or predict synergistic interactions based on the transcriptional profiles of single drugs or even drug combinations. Moreover, there is growing interest in repurposing drugs by identifying compounds that may target known or suggested ageing pathway based on the often-extensive data known about such compounds, including their three-dimensional structure bound to biological ligands, their pharmacokinetics, physical properties and known biological targets and effects^65^. Our finding support the feasibility of targeting multiple conserved ageing pathways using existing drugs to slow down biological ageing rate, an approach that, if translatable to humans, would result in dramatic medical and economic benefits^66^.

## Extended data – Tables and Figures

**Extended data Table S1.** Drug selection based on ageing pathways.

| Drugs | Proposed pathway | References |
| --- | --- | --- |
| Rapamycin | mTORC1 | 7, 11, 49 |
| Rifampicin | JNK | 24 |
| Metformin | Mitohormesis, microbial folate metabolism, AMPK | 25, 62, 63 |
| Psora-4 | K-channel blocker, Hormesis | 26 |
| Allantoin | CR mimetic | 22 |
| Torin-1 | mTORC2 | None |
| Aspirin | AMPK/DAF-16, Hormesis | 64, 65 |
| Mianserin | CR memetic | 66, 67 |
| Loxapine Succinate | Unknown | 26 |
| Juglone | ROS | 68 |
| cAMP | CR mimetic | 69 |

**Extended data Table S2.**
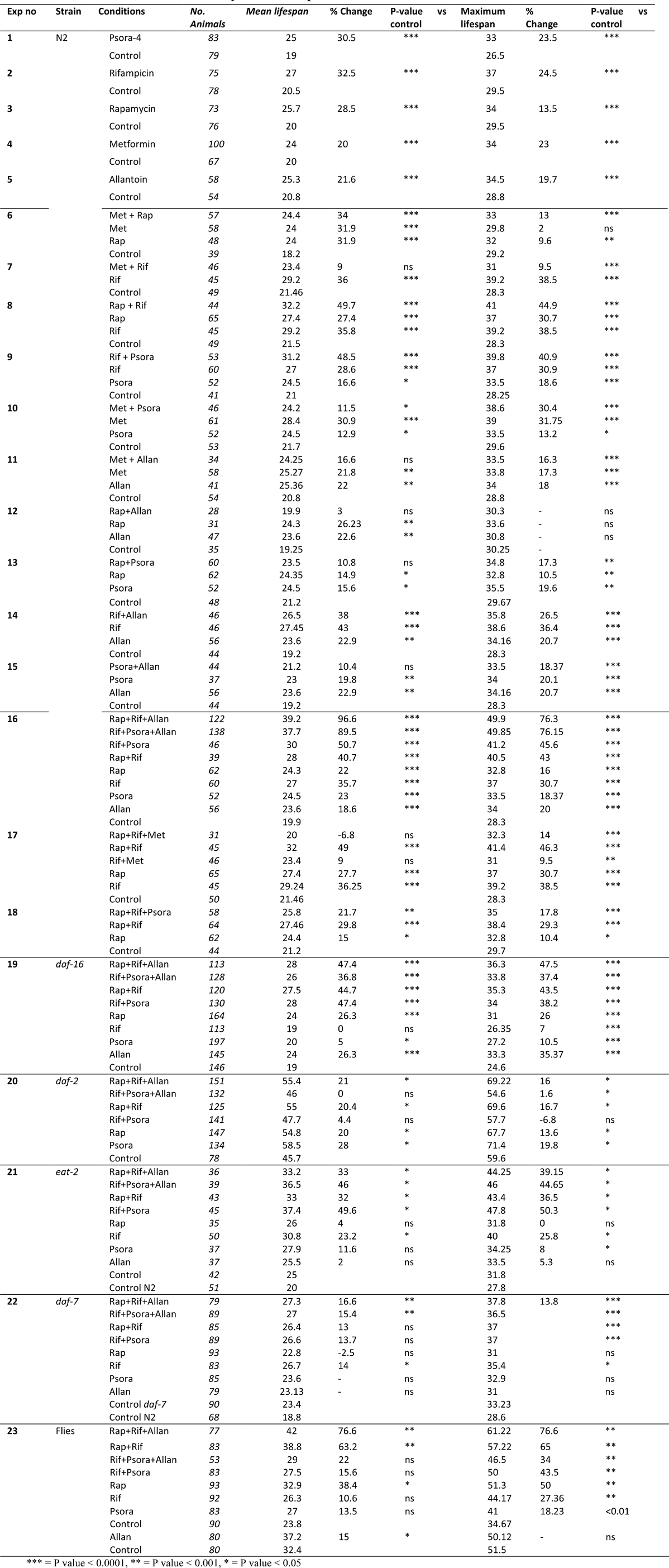
Summary of all Lifespan Studies.

***Extended data Table S3-S7 – in the attached supplementary files***

## Extended data – Figures

**Figure S1.**
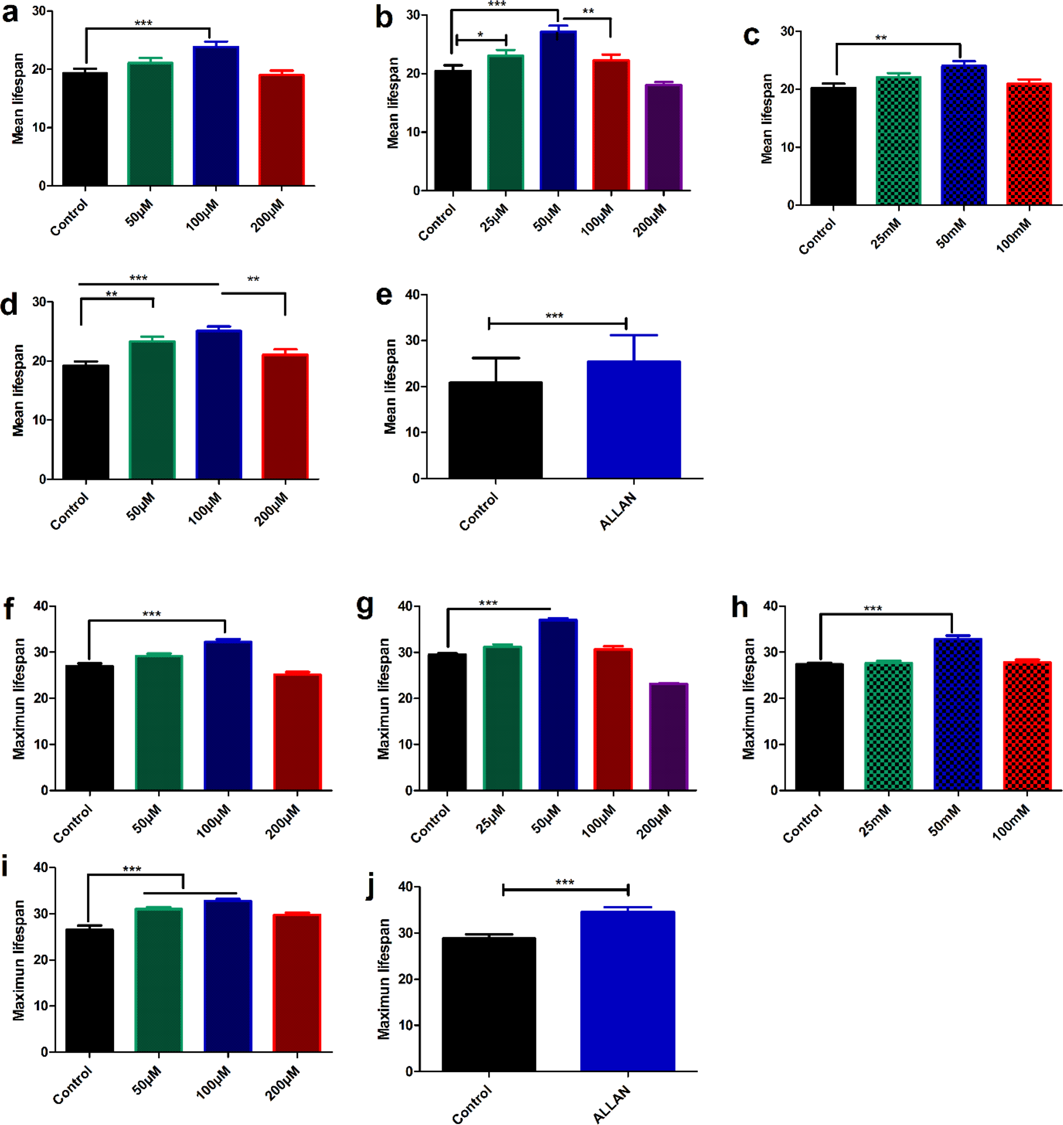
Mean and Maximum lifespan of worms treated with ranges of doses of different drugs. Drug treatment started at day 4 and continued until all worms died. Worms were transferred to fresh plate every day 3-4 days. (**a-e**) Mean and (**f-j**) Maximum lifespan of single drug treatments supplementary to figure 1. a and f – RAP, b and g – RIF, c and h – MET, d and i – PSORA, e and j – ALLAN (Mean ± SE). One-way ANOVA, Bonferroni multiple comparison ***P < 0.0001, **P < 0.001, *P < 0.01

**Figure S2.**
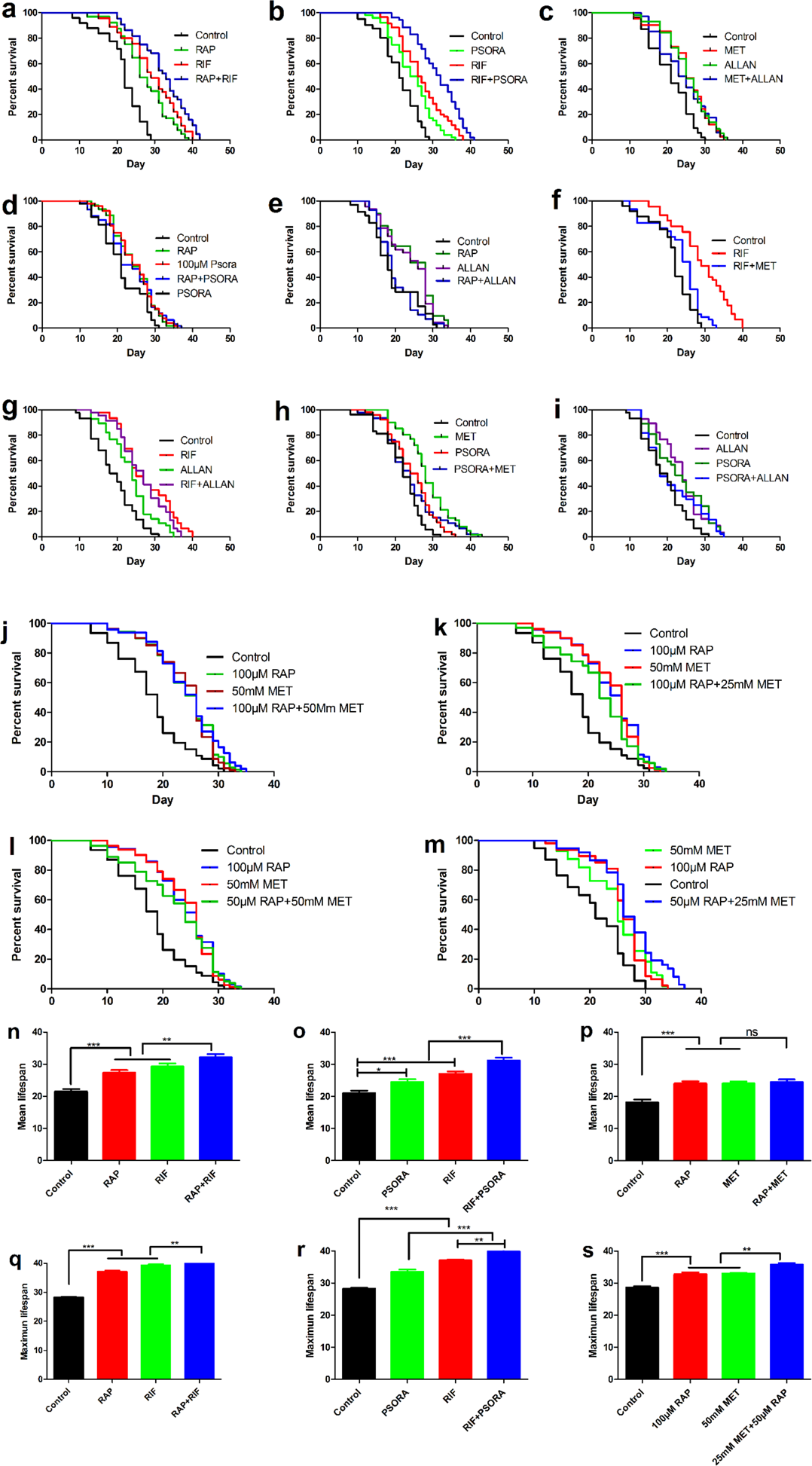
lifespan comparison of dual combinations. supplementary for figure 2. **a-i**, survival curve for all 10 possible combinations of five drugs at their respective optimal dose except RAP+MET. **j-m**, all possible combinations of RAP and MET at their optimal and half-optimal doses. **n-p**, Log-rank with proper adjustment. Mean lifespan of representative dual combinations, (Mean ± SE) **q-s**, maximum lifespan of representative dual combinations, (Mean ± SE). One-way ANOVA, Bonferroni multiple comparison ***P < 0.0001, **P < 0.001, *P < 0.01.

**Figure S3.**
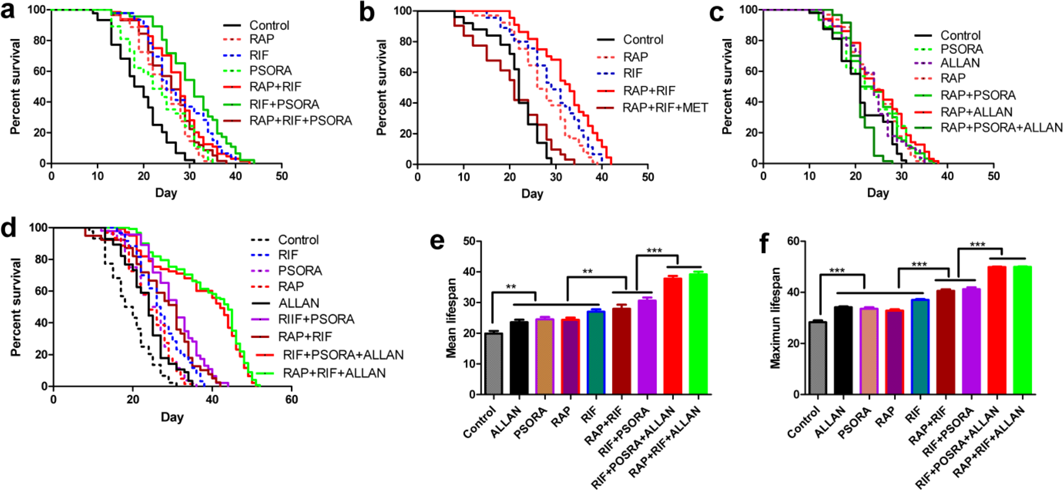
lifespan comparison of triple combinations. supplementary for figure 2. **a**, RAP+RIF+PSORA did not show further benefit compared to the dual synergistic combinations RAP+RIF and RIF+PSORA. **b**, RAP+RIF+MET is toxic compared to RAP+RIF. More than 50 worms per condition. **c**, RAP+PSORA+ALLAN did not show further benefit compared to single drug constituents. **d**, RAP+RIF+ALLAN and RIF+PSORA+ALLAN resulted in a significant lifespan extension compared to RAP+RIF and RIF+PSORA respectively. log-rank with proper adjustment. **e**, Mean and **f**, Maximum lifespan of figure d showed a monotonic increase from single drugs to double and triple combinations. One-way ANOVA, Bonferroni multiple comparison ***P<0.0001, **P<0.001, *P<0.01.

**Figure S4.**
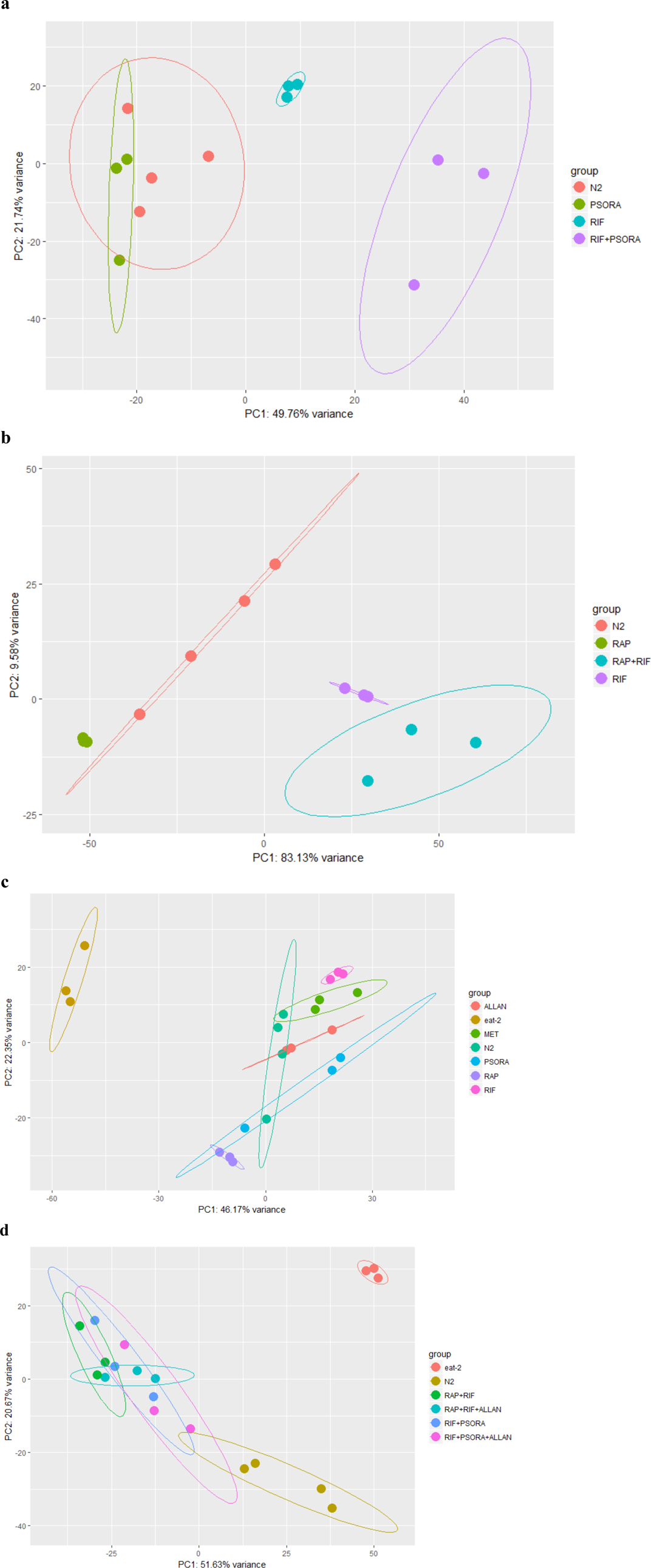
2D Principle component analysis (PCA), Transcriptome profile based on the differentially expressed genes of each drug and drug combination projected onto top two PCs. **a**, 2D PCA of PSORA, RIF and their combination showed that the combination transcriptome profile is different from single drugs. Because the single drugs and the combinations were sequenced in two different lanes, we used the N2 untreated controls of both lanes. **b**, as in **a** for RAP and RIF. **c**, similar to the 3D PCA, RAP, RIF and PSORA were well separated with 2D PCA. All single drugs were separated from *eat-2*. **d**, all synergistic combinations are well separated with N2 control and eat-2, whereas they are not separated from each other. This could show that they may have the same mechanism of synergistic lifespan extension.

**Figure S5.**
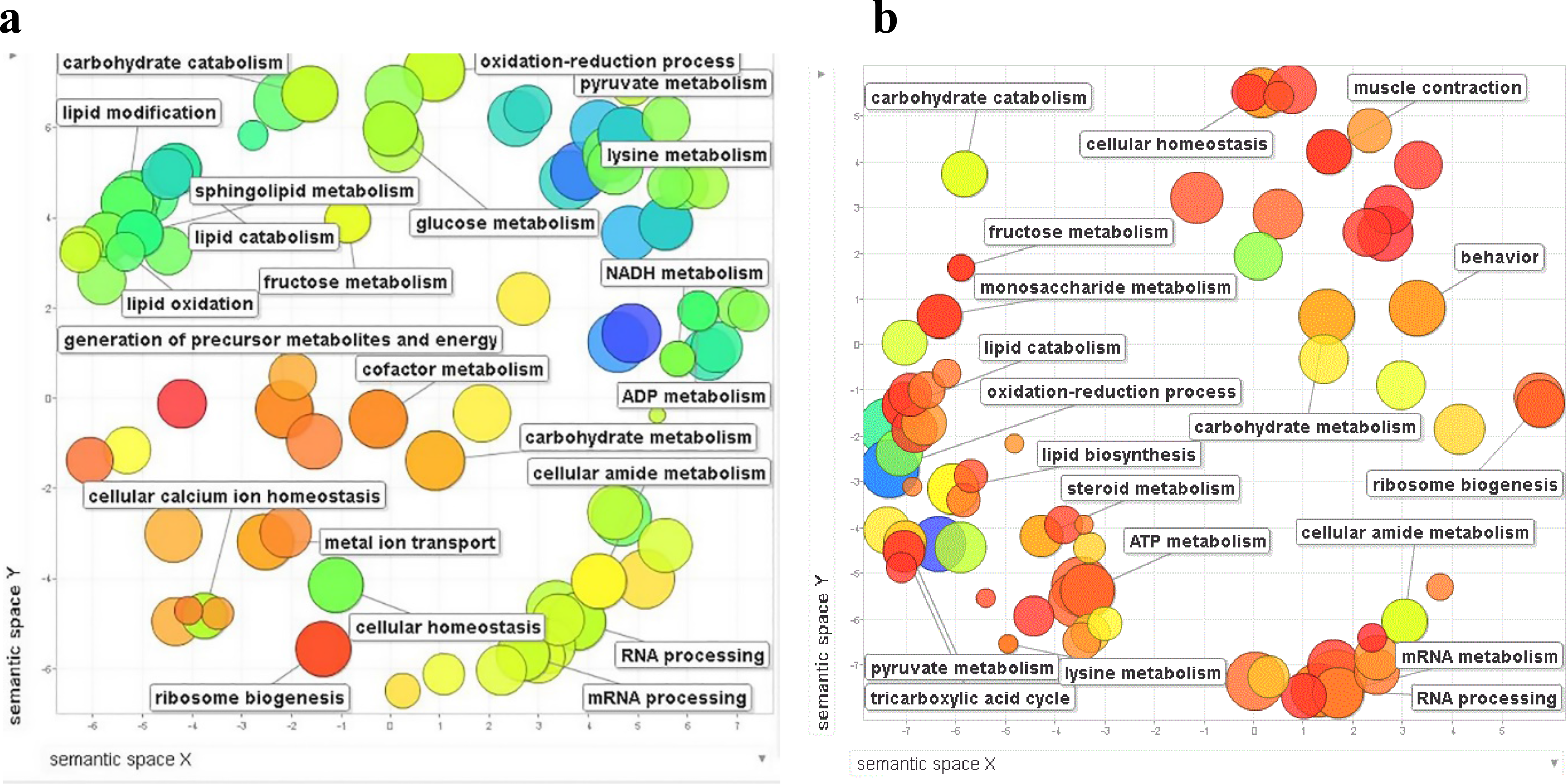
GO terms enriched by synergistic triple drug combinations determined by Cytoscape70 and summarized by Revigo71. Differentially expressed genes (|LFC| >= 1, P value < 0.05) for each conditions were used as an input for GO term analysis. both **a**, RIF+PSORA+ALLAN, and **b**, RAP+RIF+ALLAN mainly targets metabolic pathways. (P value < 0.05, term enrichment > 1.5)

**Figure S6.**
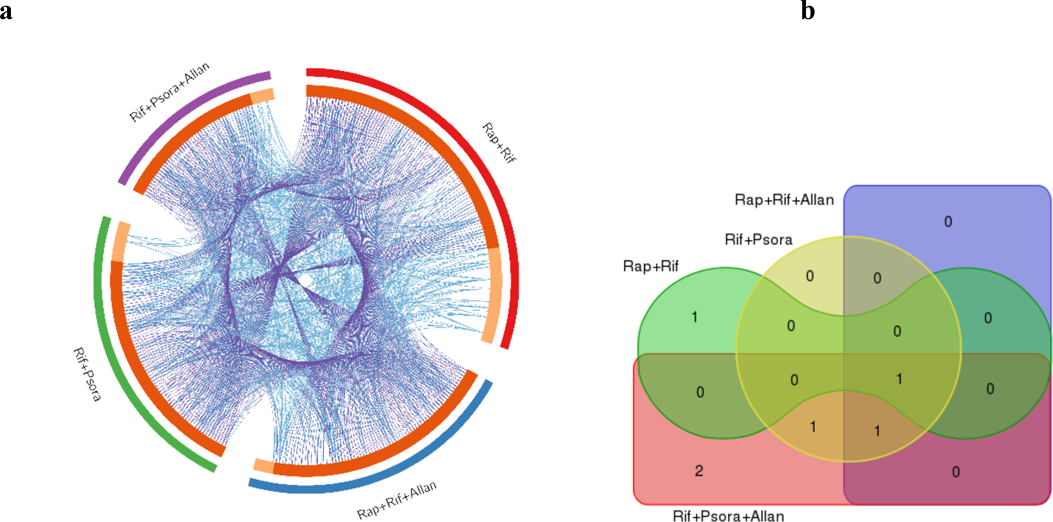
Gene signature and pathway overlap for synergistic combinations: supplementary for figure 5. **a**, DGE and GO term overlap among synergistic combinations. Purple line links identical genes whereas blue line links different gene that grouped into similar GO terms. **b**, Venn diagram of pathways enriched in synergistic combinations using *eat-2* transcriptome as a background. Only TGF-beta was commonly enriched in all synergistic combinations.

**Figure S7.**
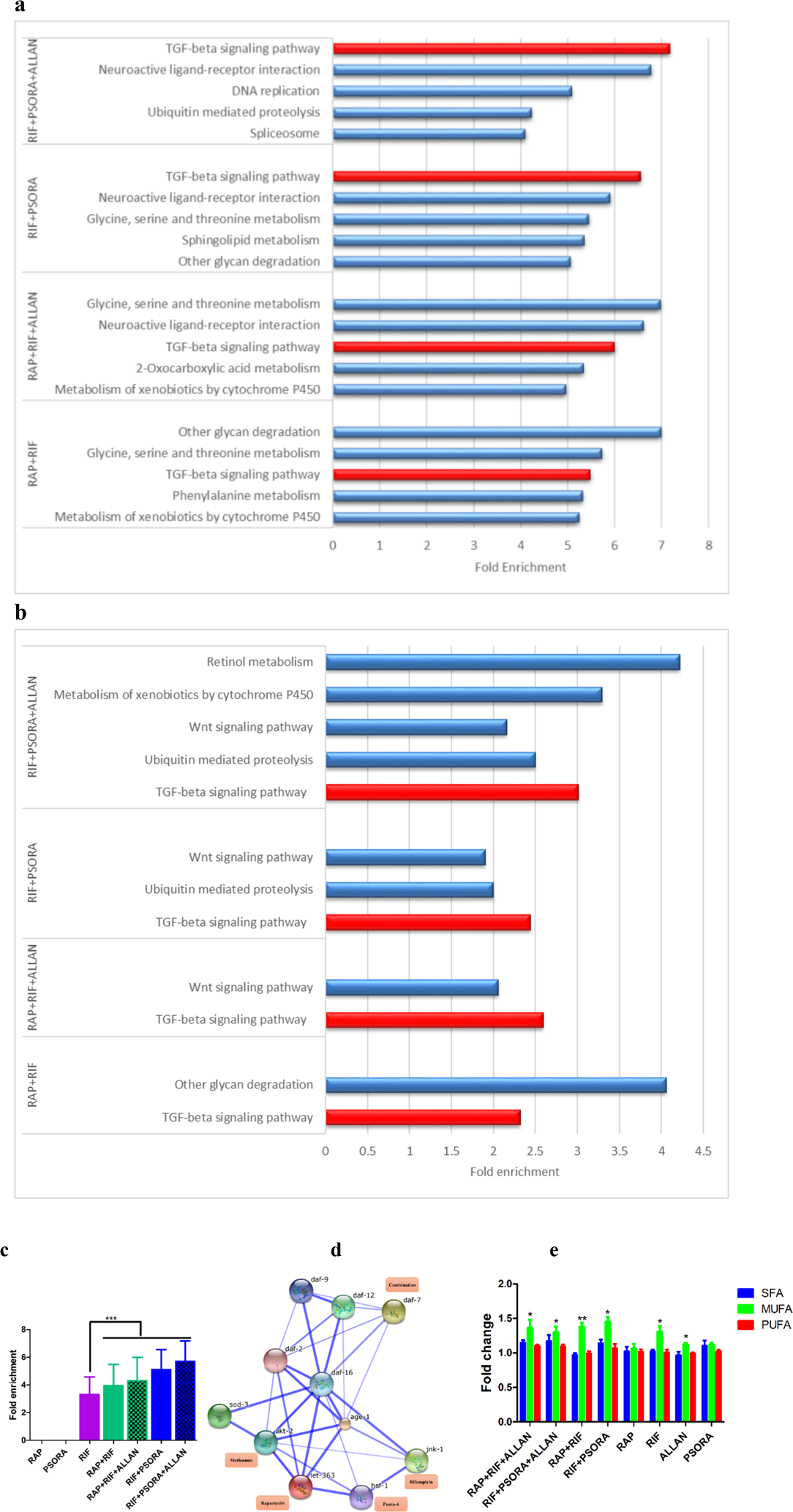
TGF-beta pathway is significantly enriched in the synergistic drug combinations differentially expressed genes. **a,** Top five pathways enriched in the down regulated genes of the four synergistic drug combinations. All synergistic combinations consistently enrich TGFβ. **b**, Pathways enriched in N2 animals treated with synergistic drug combinations relative to untreated eat-2 worms. All synergistic combinations consistently enrich TGFβ. **c**, fold enrichment of TGFβ in all single drugs and synergistic combinations. From single drugs, only RIF enriched TGFβ. All the four synergistic combinations resulted in a significant enrichment of TGFβ compared to RIF. **d**, protein-protein interaction drawn by STRING72 showed that TGFβ/*daf-7* has an interaction with *daf-2* and *daf-16* as well as the lipid metabolism regulators *daf-9* and *daf-12*. Thickness of the edge indicates strength of data support for the specific interaction. **e**, PC’s pooled based on their degree of unsaturation, an increase shown in the MUFA containing species normalized to control, 2500 worms per condition (Mean ± SD) *P<0.01, **P<0.001, ***P<0.000.

**Figure S8.**
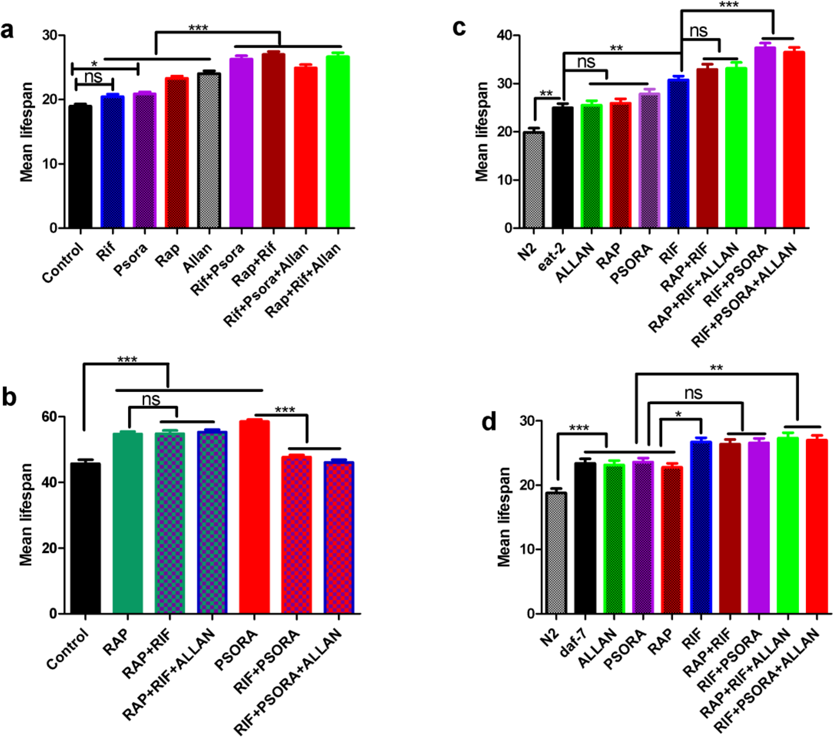
Mean lifespan on *daf-16, eat-2, daf-2* and *daf-7* mutants treated with single drugs and drug combinations. **a**, Dual combinations RAP+RIF and RIF+PSORA showed synergy in *daf-16* where as the triple combinations RAP+RIF+ALLAN and RIF+PSORA+ALLAN did not result in significant difference from their respective synergistic dual combinations. **b**, All synergistic combinations did not show synergy in *daf-2* mutant worms. **c**, Only RIF+PSORA showed synergy in *eat-2* mutants. **d**, The triple combinations showed further lifespan extension compared to single drugs but none of the four combinations show synergy in *daf-7* mutants. Supplementary for figure 4. At least 50 worms per condition. One-way ANOVA, Bonferroni multiple comparison ***P<0.0001, **P<0.001, *P<0.01.

**Figure S9.**
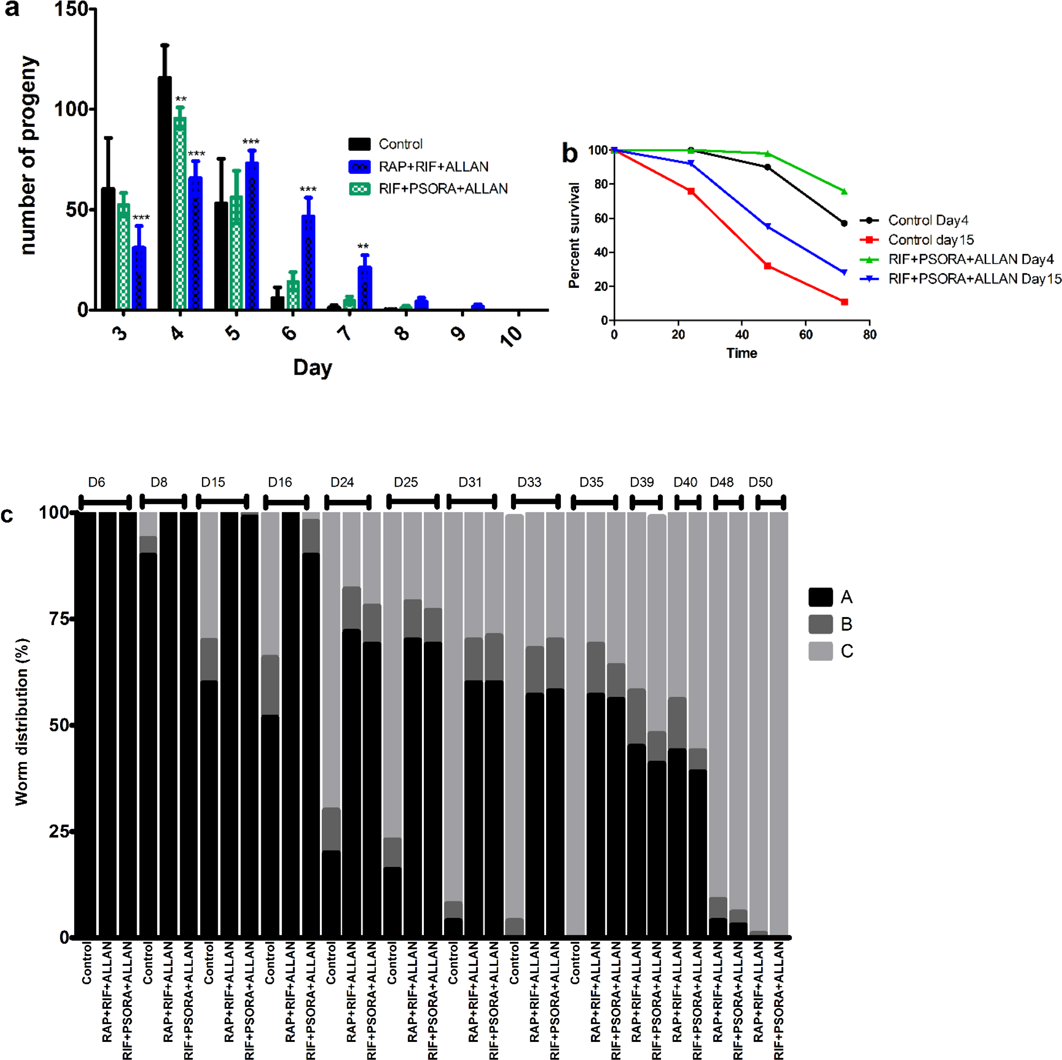
Figure S9 Health span and tradeoffs supplementary for figure 3. **a**, Drug synergies extended reproductive period. 10 worms per condition. **b**, Heat shock stress, RIF+PSORA+ALLAN improves heat shock stress resistance both at young, day 4, worms and older, day 15, worms **c**, Health span scoring at different age. At all ages from young to old drug synergy treated worms have better health span than their respective age matched controls. ***P<0.0001, **P<0.001, *P<0.01.

**Figure S10.**
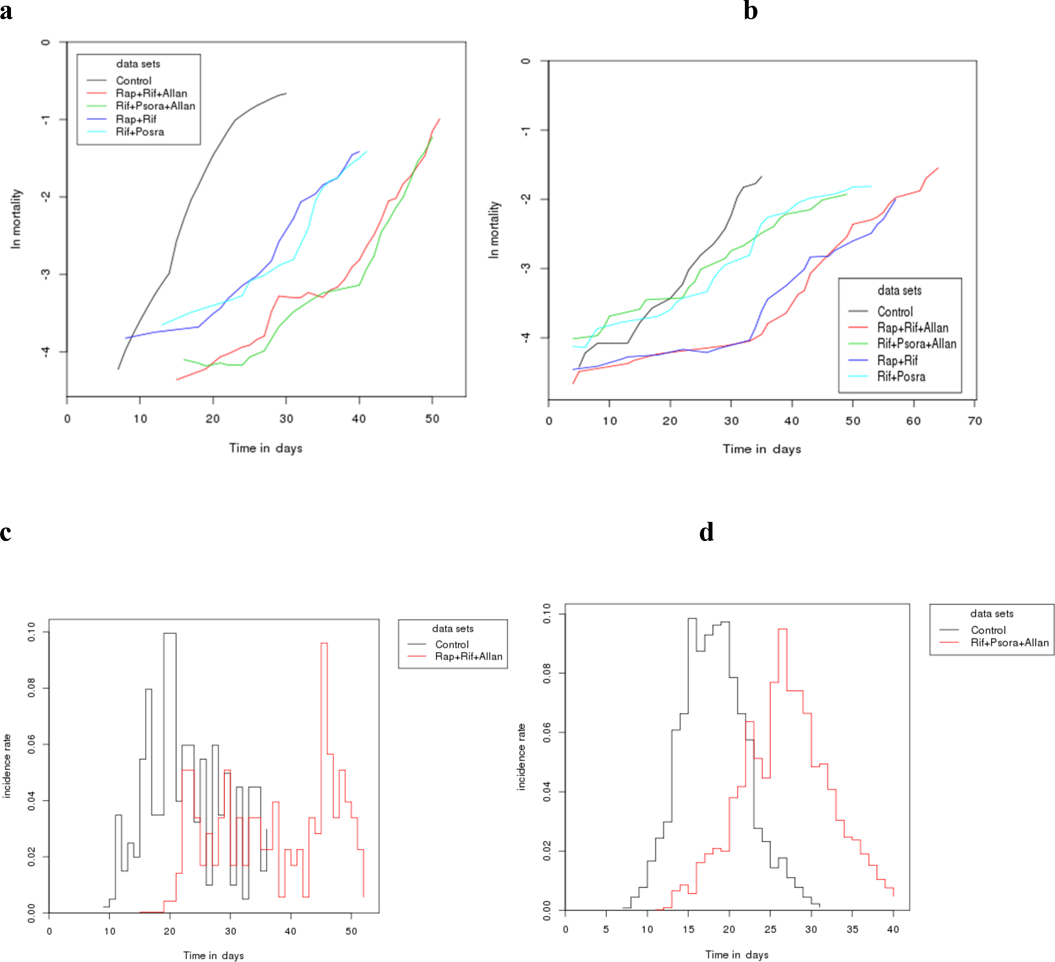
Figure S10. Drug synergy slows mortality rate and delays death incidence rate; **a**, Both the dual and triple synergistic combinations slow the rate of mortality of wild type N2 worms. **b**, The triple combinations RAP+RIF+ALLAN and RIF+PSORA+ALLAN slows mortality rate in *Drosophila Melanogaster*. **c**, RAP+RIF+ALLAN and **d**, RIF+PSORA+ALLAN delays incidence rate of death of worms.

## Supplementary Information: Supplementary Methods

### Culturing *C. elegans*

For all experiments Bristol N2 wild-type or mutant nematodes were grown and maintained on Nematode Growth Medium (NGM) agar plates at 20°C using *E. coli* OP50 bacteria as food source unless otherwise noted. After plates were poured and dried, they were sealed and stored at 4°C. *E. coli* were spotted on plates on the previous evening and allowed to dry. For compound treatments, all agar plates were prepared from the same batch of NGM agar and treatment plates were supplemented with the respective compounds or vehicle as a control. Fresh plates were prepared every week. The following worm strains were used: wild type N2, *eat-2* (DA1116), *daf-2* (CB1370), *daf-16* (CF1038), *daf-7* (CB1372). The strain CB1372 was provided by Takao Inoue. All the other strains were obtained from the Caenorhabditis Genetics Center (CGC).

### Bacterial Preparation Method

A single colony OP50 *E. coli* was picked and incubated at 37°C for 20 hours with shaking at 180 rpm in LB broth supplemented with Streptomycin (final concentration 200µg/ml). Bacterial colony forming units (CFU) were determined spectrophotometrically at 600nm wavelength, and bacterial stock was diluted to 1010, and frozen at −80°C. NGM plates were seeded with bacteria and used for lifespan assays without further incubation. Drugs or vehicles were added to the bacteria (in addition to being added to NGM) just before seeding.

### Determination of lifespan

For determination of *C. elegans* lifespan, nematodes were age synchronized by bleaching, allowed to hatch and 150-200 young adult worms per condition were transferred to 3-5 35mm culture plates. Worms were transferred to fresh plates every day until progeny production ceased, and every two to three days afterwards, until all worms had died. The number of worms that were alive was determined every day, and dead worms were removed from the plate. Worms were considered dead when no touch-provoked movement was observed. Animals that crawled off the plate, exhibited extruded internal organs or displayed an egg laying defective with subsequent hatching of larvae inside the mother were censored. All lifespan assays were blinded and repeated at least two times. Graph pad Prism 5.0 software was used to calculate mean adult lifespan and to perform statistical analysis of significance using the log-rank test (Mantel-Cox).

### Compound identification and validation

#### Primary Screening

First, we identified well-characterized ageing regulatory pathways (Extended data Table-1). Next, we identified drugs and drug-like molecules that are known to target the selected pathways and to extend lifespan in a common ageing model organisms (worm, fly or mice). Some compounds that had been reported to target known ageing pathways and had not yet been experimentally validated as lifespan-extending were also included. This approached culminated in the identification of 11 candidate drugs (Extended data Table-1). Primary efficacy was confirmed in terms of lifespan in *C. elegans*. Initial screening of compounds for their effect on lifespan of wild type N2 nematodes were conducted using standardized condition in the presence of antibiotics (200mg/ml streptomycin). Of these 11 drugs and drug-like molecules, only five resulted in reproducible lifespan extension in our lab (Fig. 1). For those compounds that passed the primary screening, dose-response assays were carried out to establish the ideal dose and maximum effect size.

### Resistance to oxidative stress

Age-synchronized adult worms were transferred to a fresh NGM plate containing drug or vehicle. 5-fluoro-2’-deoxyuridine (FUdR) was also added to prevent egg hatching. After 48 hours of treatment, worms were transferred to plates with 20mM paraquat (methyl viologen dichloride hydrate, Sigma Aldrich) to induce oxidative stress in the worms. Thereafter, dead worms were counted every day until all of the worms were dead.

### Heat shock stress resistance

Age-synchronized adult worms were picked from a NGM plate and transferred to a fresh NGM plate containing drug or vehicle. After 48 hours of treatment, worms were transferred to 37°C incubator for 2 hours to induce heat stress. After heat shock worms were transferred back to 20°C. Survival rates were determined after 24 hours, 48 hours and 72 hours of recovery and compared between the control and drug-treated worm groups.

### Fertility assay

Age-synchronized day 3 worms were picked from NGM plate and transferred to fresh NGM plate containing drug or vehicle. A single worm was used per plate with 10 plates per condition. Worms were transferred to a fresh NGM plate containing drug or vehicle every day until they ceased egg laying. Eggs produced by each single worm were incubated at 20°C for 48 hours, and the number of live progeny produced was recorded on each day.

### Development by size

Age-synchronized L2 worms were transferred to fresh NGM plates containing drug or vehicle. We took images of worms every day, starting from day 4, the first day of drug treatment for all experiments, up to day 11. Length of worms was determined using the free curve tool provided by the Leica Application Suite software (Leica Microsystems). We compared the mean size of worms under different treatments with the size of untreated control worms in each day using t-test.

### Respiration rate

Worms were treated with drug or vehicle for 24 hours. We transfer worms to 96 well assay plates, nine worms per plate at least eight wells per condition. Respiration rates were determined using Seahorse XF96 respirometer following standard protocol^1^.

### Statistical methods and synergy definition

Survival analysis for all single drugs and all dual and triple combinations were determined by multiple comparison tests using log-rank test. Statistical analyses of mean and maximum lifespan difference were determined by one-way ANOVA, Bonferroni multiple comparison test to compare all pairs of columns. Furthermore maximum lifespan difference were compared using Wang-Allison Score test^2^. Allantoin mean lifespan difference was determined by T-test. To determine whether a drug combination is synergistic first we determined the percent lifespan extension by each individual drug and compared this effect to the effect of their combination. We consider a combination of drugs to be synergistic if the lifespan effect under combinational treatment is significantly greater than the lifespan effect of the more efficacious of the individual drugs. For triple drug combinations, we test the combined effects against the sum of the effects of the best pair out of the three drugs.

### Mortality

We experimentally determined the mortality rate of one of our most powerful interventions, RIF+PSORA+ALLAN, using a large cohort of over 1,000 worms per condition. Dead worms were counted every day until all worms were dead. For RAP+RIF+ALLAN the combined lifespan data was imported to Survcurv for mortality analysis and maximum lifespan comparison^3,4^. We used the Gompertz survival model to determine the initial mortality rate and mortality rate doubling time using the following equation.

M(t) = ae^bt^

MRD = ln2/b, where

a = baseline hazard rate

b = rate if exponential increase in mortality

t = time

M(t) = mortality rate

MRD = mortality rate doubling time

### RNA Extraction and RNAseq

Freshly prepared bacteria (OP50 *E. coli*) were spotted on 94 cm NGM agar plates on the previous evening and allowed to dry. Synchronized, young adult worms were transferred to fresh plates. For maintaining synchronized populations, FUdR was added to NGM media. After two days of treatment with drug or vehicle adult worms were washed off the plates to 15 ml tubes and washed several times until a clear solution, that indicates bacteria is completely removed, was obtained. Clean worm pellets were then frozen and used for RNA extraction. Total RNA was isolated using Qiagen RNAeasy micro kit (Qiagen, Hilden, Germany) following the standard protocol. Afterwards, RNA was quantified photometrically with NanoDrop 2000 and stored at −80°C until use. The integrity of total RNA was measured by Agilent Bioanalyzer 2100. For library preparation an amount of 2µg of total RNA per sample was processed using Illumina’s RNA Sample Prep Kit following the manufacturer’s instruction (Illumina, San Diego, CA, USA). All single drugs were processed together and multiplexed onto one lane to minimize batch effect. The different drug combinations were processed into two multiplexed lanes. In total, we ran three lanes in parallel with 20 samples in each and we had untreated N2 controls in each lane. Libraries were sequenced using Illumina HiSeq4000 sequencing platform (Illumina, San Diego, CA, USA) in a paired end read approach at a read length of 150 nucleotides. Sequence data was extracted in FastQ format. The RNAseq reads from each sample were mapped to the reference *C. elegans* transcriptome (WBcel235) with *kallisto* (v0.43.0)^5^ using 100 bootstrap samples and sequence based bias correction. The estimated counts were imported from kallisto to the R environment (v3.3.2) and summarized to gene-level using the *tximport* package (v1.2.0)^6^. The *DESeq2* package (v1.14.1)^7^ was used to identify differentially expressed genes (DEGs) in all our analysis (correcting for batch effects, when applicable) for a significance threshold α of 0.05 after correcting for multiple hypothesis testing through Independent Hypothesis Weighting (IHW package v1.2.0)^8^. The PCA plot were graphed by the *pcaExplorer* package (v2.0.0) on DESeq2 ‘rlog’ transformed data.

### Analysis with known lifespan-extending genes

All 569 *Caenorhabditis elegans* genes that extend lifespan when intervened upon were downloaded from GenAge (build 18)^5^ and of those, 512 were verified to be present in our genes expression datasets, and this subset of genes were used exclusively in order to graph the respective heatmap. The heatmaps only used gene expression counts (after DESeq2 normalization and replacement of outliers) for this subset of genes. All the samples of each condition were aggregated by taking the medium values of gene expression counts.

### Pathway Analysis

KEGG pathways and GO terms enriched by each drug and drug combination were determined by DAVID^9^ and Metascape^10^. To determine pathways enriched in combinations compared to single drugs, we used the differentially expressed genes by single drugs as a background control. To determine the pathways enriched by the synergistic drug combinations compared to *eat-2* we used the genes differentially expressed in untreated *eat-2* vs untreated N2 worms as a background. Venn diagram of pathways done online using (http://bioinformatics.psb.ugent.be/webtools/Venn/)

### Lipid extraction and Mass spectrometry

#### Lipid extraction

– young adult day 4 worms, 2,500 per condition, were treated with drug or vehicle for 48 hours. Nematodes were collected, washed with M9 buffer and transferred to 2-mL polypropylene tubes containing 250µl lysis buffer (20mM Tris-HCl pH 7.4, 100mM NaCl, 0.5mM EDTA, 5% glycerol) and left on ice for 15 minutes followed by homogenization using a Bead beater maintained at 4°C. Lipids extraction from the lysed samples was carried out by Folch’s extraction^11^. In order to minimize oxidation during and after extraction, 0.5% butylated hydroxytoluene was added to the organic solvents. The upper phase was removed, and to the lower phase a theoretical upper phase was added, vortexed and centrifuged at 3000rpm for 5 minutes. The lower phase was then transferred to a tube, evaporated and dried under vacuum to avoid lipid oxidation. For quantification purposes, the organic phase was spiked with a labeled internal standard corresponding to each lipid class during the single phase extraction so as to control lipid-class dependent differences in extraction and ionization^12^. Standards used were PC-34:0, PE-28:0, SM-30:1, Cer-35:1, DAG-24:0 and TAG-51:0. Subsequently, the samples were evaporated under a speed-valco (Thermo Savant, Milford, USA) and reconstituted in 50µl methanol and left at −80°C until further analysis.

#### Data acquisition

An Agilent 1260-Ultra Performance Liquid chromatography (UPLC) coupled to Triple Quad Mass spectrometer (Agilent 6490) with dynamic multiple reaction monitoring (dMRM) was used for lipid quantification. The UPLC system was equipped with a Waters ACQUITY BEH C18column (1.0 × 100 mm) to separate the molecular species using gradient elution. Solvent A was acetonitrile/H2O (60:40) with 10mM ammonium formate and 1% NH_4_OH, while solvent B was isopropanol/acetonitrile (90:10) containing 10mM ammonium formate and 1% NH_4_OH. The flow rate was 0.13mL/min and the column temperature 60°C. Solvent B was set at 40% at injection and increased linearly to 100% in 14 minutes, retained at this value for 3 minutes, decreased back to 40% in one minute and then retained there till the end of the gradient by 20 minutes. The eluent was directed to the ESI source of the mass spectrometer operated in the positive ion mode. The MS conditions were as follows. For ESI: gas temperature, 300°C; gas flow, 10 l/minutes; sheath gas temperature, 350°C; sheath gas flow, 8 l/minutes; and capillary voltage, 3,500 V. For APCI: gas temperature, 300°C; vaporizer, 450°C; gas flow, 5 l/minutes; capillary voltage, 4,000 V; and corona current, 4 µA. Data processing, including peak smoothing and integration of areas under the curves for each ion measured, was performed using the MassHunter Quantification Software (Agilent).

### Fly stocks, husbandry and lifespan study

Wild type fruit flies, *Drosophila melanogaster* Oregon-R, were maintained and experiments were conducted at 28oC because fly lifespan at this temperature allows for more rapid lifespan studies. Flies were maintained on food containing 6g Bacto agar, 114g glucose, 56g cornmeal, 25g Brewer’s yeast and 20ml of 10% Nipagin in 1L final volume. Newly enclosed flies were raised in standard bottle and transferred to new food bottles daily at Day 1 of age. Male flies were sorted into small vials (12 flies per vial) on to the food tube with drugs applied. Lifespan counts were recorded five times a week for death and censors. Flies were transferred to fresh food tubes with corresponding drug treatments once every three weeks.

